# A soybean rust effector protease suppresses host immunity and cleaves a 3-deoxy-7-phosphoheptulonate synthase

**DOI:** 10.1101/2023.09.07.556260

**Authors:** Aline Sartor Chicowski, Mingsheng Qi, Haris Variz, Melissa Bredow, Christian Montes, Francesco Caiazza, Haili Dong, Alexandra Margets, Joffrey Mejias, Justin W. Walley, Charles S. Craik, Kerry F. Pedley, Kyaw Aung, Roger Innes, Steven A. Whitham

**Affiliations:** Department of Plant Pathology, Entomology and Microbiology, Iowa State University, Ames, Iowa, USA; Department of Genetics, Development, and Cell Biology, Iowa State University, Ames, Iowa, USA; Department of Pharmaceutical Chemistry, University of California, San Francisco, USA; Department of Biology, Indiana University, Bloomington, Indiana, USA; PHIM Plant Health Institute, University of Montpellier, INRAE, CIRAD, Institut Agro, IRD, Montpellier, France; United States Department of Agriculture-Agricultural Research Service (USDA-ARS), Foreign Disease-Weed Science Research Unit (FDWSRU), Fort Detrick, MD, USA

**Keywords:** Plant immunity, soybean rust, plant defenses suppression, rust effectors, effector protease

## Abstract

The devastating soybean rust (SBR) pathogen, *Phakopsora pachyrhizi*, encodes many secreted proteins, but only two have been functionally characterized for their roles in rust virulence. Here, we demonstrate that transient expression of *P. pachyrhizi* effector candidate 15 (*Pp*EC15), an aspartic protease, leads to enhanced bacterial growth *in planta*, suppression of callose deposition, reduced expression of plant defense-related marker genes and suppression of pathogen-associated molecular pattern (PAMP)-induced reactive oxygen species (ROS). Stable expression of *Pp*EC15 in soybean suppresses PAMP-induced ROS production and enhances bacterial growth, indicating that, collectively, *Pp*EC15 suppresses host and non-host innate immune responses. Yeast-two-hybrid and proximity labeling identified putative *Pp*EC15 interacting partners including a peptide-chain release factor (PCRF), a NAC83 (NAM, ATAF, and CUC) transcription factor, and a DAHP (3-deoxy-7-phosphoheptulonate) synthase. We further show that *Pp*EC15 can cleave DAHP but does not cleave PCRF or NAC83. Virus-induced gene silencing of NAC83, PCRF and DAHP altered PAMP-induced ROS production and salicylic acid production, indicating that these proteins may be involved in immune signaling. Collectively, our data show that *Pp*EC15 is conserved across *P. pachyrhizi* isolates and other economically important rust species and is involved in the suppression of plant basal defense responses. Understanding the role of *Pp*EC15 in *P. pachyrhizi* virulence will provide a foundation for designing targeted intervention strategies to generate rust-resistant crops.

## Introduction

Rust fungi, in the order Pucciniales, are among the most economically important and devastating pathogens of crop plants, causing yield losses in the order of billions of dollars annually (Helfer, 2014). Soybean rust (SBR), caused by the highly adaptive and obligate biotrophic fungus *Phakopsora pachyrhizi*, is a devastating foliar disease affecting soybean. Yield losses caused by SBR can be as high as 90%, and even low disease incidence can negatively impact yield (Godoy et al., 2016). So far, there is no soybean cultivar available that is fully resistant to all *P. pachyrhizi* isolates present in soybean-growing areas, resulting in heavy use of fungicides to control *P. pachyrhizi* outbreaks (Godoy et al. 2016). Because of the complexity of the pathosystem, and the lack of durable sources of resistance, determining the underlying molecular and genetic factors that contribute to SBR virulence is an essential step toward developing plants resistant to *P. pachyrhizi*.

In general, the establishment of plant diseases caused by biotrophic pathogens requires the deployment of an arsenal of effectors that manipulate the structure and function of plant cells in favor of disease progression (Bentham et al., 2020). Pathogen-secreted effectors can function in the plant apoplast or can be translocated to the host cytoplasm after secretion (Selin et al., 2016). To counteract these measures, plants recognize pathogens through an innate immune system that monitors pathogen-associated molecules either outside or inside the plant cell by relying on two layers of immunity with shared signaling components, ultimately resulting in many overlapping downstream outputs (Yuan et al., 2021). The first line of defense is known as pattern-triggered immunity (PTI) and is mediated by the recognition of pathogen molecules conserved within a class of microbes, known as pathogen-associated molecular patterns (PAMPs), by plant transmembrane pattern-recognition receptors (PRRs) (Segonzac and Zipfel, 2011). PAMP recognition initiates an intracellular signaling cascade leading to broad-spectrum disease resistance and includes the generation of apoplastic reactive oxygen species (ROS), influx of extracellular calcium (Ca^2+^), upregulation of defense-related genes, and the accumulation of phytohormones (Chang et al., 2022). The second layer of defense responses is mediated by intracellular nucleotide-binding leucine-rich repeat (NLR) proteins that recognize pathogen effector proteins leading to a more robust immune response known as effector-triggered immunity (ETI), often associated with a form of localized cell death known as the hypersensitive response (HR) (Duxbury et al., 2021).

Effectors can manipulate different cellular processes to promote infection and disease. These proteins can interact with multiple host targets and move between various cellular components to disrupt normal cellular processes, thereby interfering with host defense responses (Nirmala et al., 2011; Sperschneider et al., 2017, He at al., 2020). For example, some pathogen effectors have been shown to be active proteases that can alter or cleave defense-related host proteins for their benefit (Chandrasekaran et al. 2016; Shao et al. 2003). Numerous proteases have also been shown to promote fungal development, facilitate the formation of infection structures, and degrade host cell walls (Chandrasekaran et al., 2016). Given the role of proteases in the infection strategies of other fungal pathogens, it is likely that these proteins play an important role in eliciting or modulating host immune responses in rusts (Jashni et al., 2015). However, little is known about their specific roles in rust virulence.

*P. pachyrhizi* encodes many secreted proteins (Link et al., 2014; Kunjeti et al., 2016; de Carvalho et al., 2017; Elmore et al., 2020), but only two have been functionally characterized for their roles in *P. pachyrhizi* virulence (Qi et al., 2016; Bueno et al., 2022). Previously, a large-scale screen to identify putative *P. pachyrhizi* effectors capable of suppressing or activating plant immunity was conducted by our group, in which we identified a *P. pachyrhizi* predicted effector protease named *Pp*EC15 (Qi et al., 2018). To better understand the roles of *Pp*EC15 in *P. pachyrhizi* virulence, we functionally characterized it. Our data show that *Pp*EC15 suppresses plant basal defense responses in heterologous systems and soybean, indicating that it may be a *bona fide* virulence effector. *Pp*EC15 is a functional protease, although this activity is not required for all of its virulence functions. We additionally identified three *Pp*EC15 interacting soybean proteins that modulate plant defense responses, including a peptide-chain release factor (PCRF), a NAC83 (NAM, ATAF, and CUC) transcription factor, and a DAHP (3-deoxy-7-phosphoheptulonate) synthase. We further show that *Pp*EC15 cleaves DAHP but does not cleave PCRF or NAC83. Silencing of the targeted soybean proteins altered PAMP-elicited ROS production and salicylic acid (SA) production, indicating that these proteins might be involved in soybean immunity. Our data suggests that *Pp*EC15 is a multifunctional effector that can cleave one of its interacting proteins possibly to impair its activity or function in immune signaling. Together, our data uncover a novel function for a *Pp*EC and key components of soybean immunity. Understanding the functions of effectors in the infection process is an important step in uncovering *P. pachyrhizi* infection strategies and can aid in the development of effective and long-lasting strategies to enhance SBR resistance in soybean.

## Results

### Sequence analyses of *Pp*EC15

*Pp*EC15 was previously identified as a haustoria-expressed secreted protein of *P. pachyrhizi* (Link et al., 2014). Sequence analysis suggested that *Pp*EC15 was an aspartic protease, and in a large-scale screen of 82 *Pp*ECs for effector-like functions, it was one of 17 *Pp*ECs that suppressed plant basal immune responses (Qi et al., 2018). Sequence alignment of *Pp*EC15 with two known aspartic proteases, saccharopepsin from *Puccinia graminis* f. sp. *tritici* and aspartic peptidase A1 from *Trametes versicolor*, showed that they share conserved aspartyl protease family A1A domains (yellow colored) (Figure 1A). All contain predicted amino (N)-terminal signal peptides (orange colored), suggesting they may all be secreted proteins. Only *Pp*EC15 contains a predicted nuclear localization signal (NLS) (green colored), consisting of a continuous Arg/Lys core sequence, which is absent in the other two homologs, and an Arg/Lys flanking sequence, which is present in the other two homologs (Figure 1A). *Pp*EC15 has two predicted catalytic motifs and a catalytic flap site (Figure 1, A-C). Sequence analysis shows that *Pp*EC15 is conserved among eleven *P. pachyrhizi* isolates (designated as TW, IN, CO, PG, AU, HW, BZ, BX, K8108, MG2006, and PPUFV02) from four different continents (Figure 1B). Moreover, *Pp*EC15 has a high level of conservation with many other proteases from different fungal pathogen species, including *Melampsora larici*, *Rhodotorula graminis*, *Cryptococcus* species and *Puccinia sp. avenae*.

**Figure 1.**
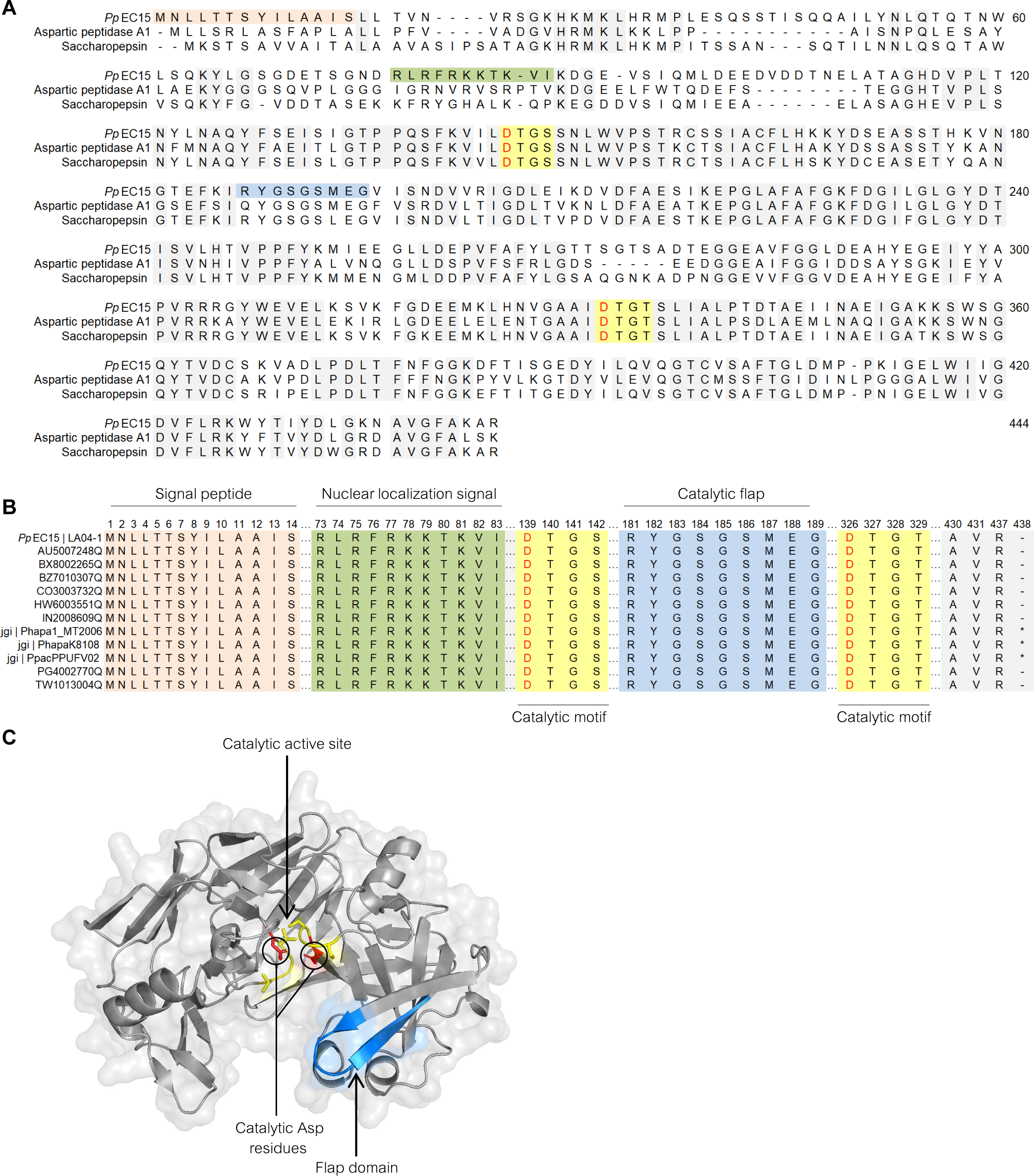
Domain analysis and structural modeling of *Pp*EC15. **A.** Sequence alignment of *Pp*EC15 (GenBank: KAI8454851.1) with saccharopepsin from *Puccinia graminis* f. sp. *tritici* (NCBI reference: XP_003320537.1) and aspartic peptidase A1 from *Trametes versicolor* (NCBI reference: XP_008035722.1). *Pp*EC15 has a predicted signal peptide (orange), a nuclear localization signal (NLS) consisting of a continuous Arg/Lys core sequence (green), two predicted catalytic motifs (yellow), catalytic Asp residues (red text), and a catalytic site flap (blue). **B.** Sequence alignment of *Pp*EC15 from 11 diverse *P. pachyrhizi* isolates. **C.** *Pp*EC15 structural modeling using the Phyre2.0 algorithm using *Saccharomyces cerevisiae* Proteinase A (PDB, 1dpj) as a template with 100% confidence score and 95% coverage. Aspartic protease domains are identified in yellow, the active site flap in blue, and the catalytic residues are in red.

### *Pp*EC15 possesses *in vitro* protease activity

To study the domain functions, three *Pp*EC15 mutants were constructed: *Pp*EC15nls1, in which all Arg/Lys in the predicted NLS were substituted with Ala (ALAFAAATAVI), *Pp*EC15nls2, in which only Arg/Lys in the core NLS sequence were substituted with Ala (RLRFAAATKVI), and a catalytic motif double mutant (*Pp*EC15dm) where the catalytic Asp residues of both motifs were substituted with Ala (DTGS-ATGS, DTGT-ATGT) (Figure 2A). To validate the predicted protease activity of *Pp*EC15, recombinantly-expressed wild-type *Pp*EC15 (*Pp*EC15wt) and the catalytic motif double mutant (*Pp*EC15dm), were incubated with casein, a generic substrate of proteases. The products of this reaction were visualized by SDS-PAGE, which showed that *Pp*EC15wt cleaved casein into two products of approximately 22 kDa and 8 kDa (Figure 2B). In contrast, *Pp*EC15dm was unable to cleave the substrate. These results indicate that *Pp*EC15 is an endopeptidase that may recognize a specific cleavage motif.

**Figure 2.**
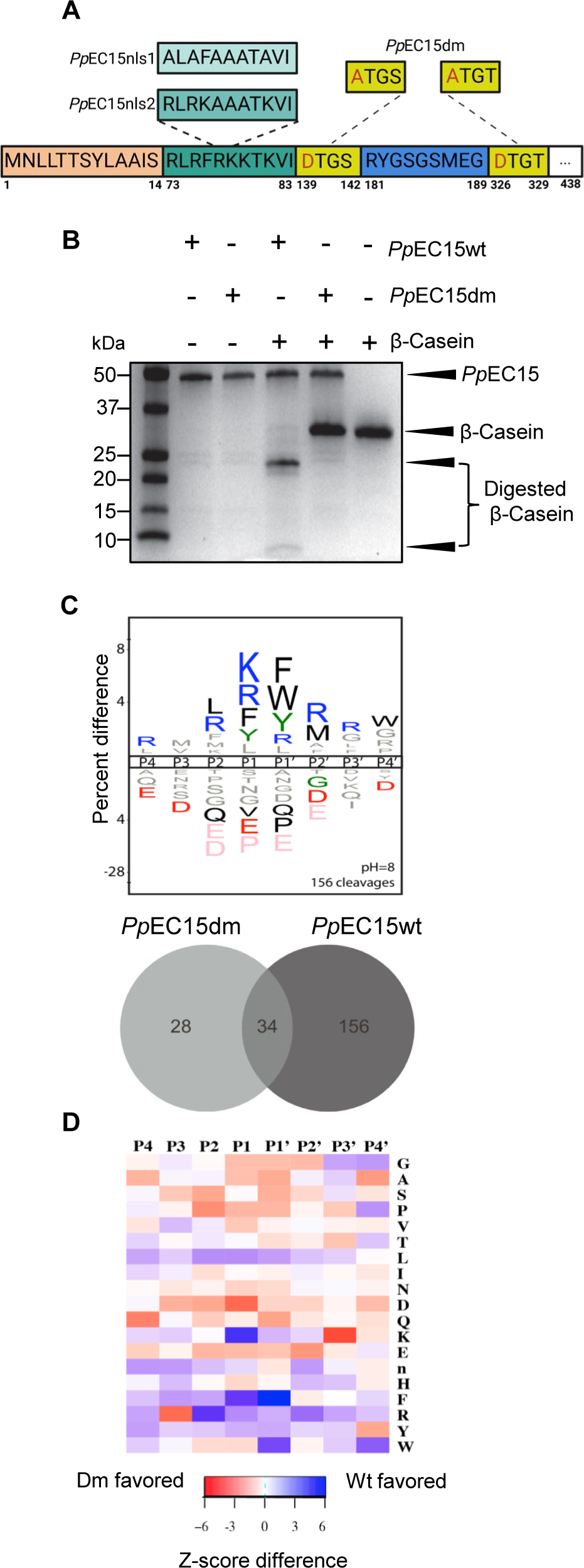
*Pp*EC15 has aspartyl protease activity. **A.** Diagrammatic representation of wild-type *Pp*EC15 and mutants generated for this study: *Pp*EC15nls1 (Ala substitutions in all the Arg/Lys residues in the nuclear localization signal (NLS)), *Pp*EC15nls2 (Ala substitutions in the core sequence of the NLS), and *Pp*EC15dm (Ala substitutions in the catalytic Asp residue). **B.** *Pp*EC15-casein cleavage assay shows that *Pp*EC15wt cleaves the model substrate β-casein into two bands consisting of 22 kDa and 8 kDa approximately, while mutations in both catalytic motifs (*Pp*EC15dm) abolished casein cleavage. **C.** Motif analysis of the 156 unique cleavages in *Pp*EC15wt samples after subtracting background from *Pp*EC15dm cleavages and quantification of the unique cleavages in the *Pp*EC15wt and *Pp*EC15dm samples in the cumulative time course assay based on the MSP-MS library of 228 14mer peptides. Residues above the line are favored; residues below the line are disfavored. Statistically significant residues (P <0.05) are colored by physicochemical properties (black: hydrophobic; blue: basic; red: acidic). Residues in gray were not statistically significant, but they were observed, and residues in pink are absolutely disfavored (never observed in a cleavage). Letter height corresponds to the fold enrichment value. **D.** Differences in Z-scores at the P4-P4’ positions are represented as a heatmap to highlight residues that are favored in *Pp*EC15wt-specific cleavages (“n” is Nle, a proxy for Met).

### *Pp*EC15 peptide cleavage specificity is consistent with aspartyl peptidase activity

To further investigate the cleavage specificity of *Pp*EC15, a time-course peptide cleavage assay was performed utilizing a library of 228 14-mer synthetic peptide substrates (O’Donoghue et al., 2015) incubated with *Pp*EC15wt or *Pp*EC15dm. Peptide cleavage products from the Multiplex Substrate Profiling by Mass Spectrometry (MSP-MS) library were identified through Liquid Chromatography with tandem mass spectrometry (LC-MS/MS). *Pp*EC15wt generated a total of 270 cleavages during the time course of 60 min, 240 min, and 1200 min. The profile of substrate specificity was extrapolated from the cumulative list of 270 cleavages observed across the entire time course and is reported using iceLogo representation (Supplemental Figure S1A). IceLogo was used to visualize the fold enrichment and de-enrichment of amino acids flanking the cleavage site at (P1-P1’) and considers both cleaved and uncleaved positions in the peptide library. The specificity profile suggests that *Pp*EC15 is an active endopeptidase, with a strong preference for positively charged amino acids at position 1 (P1) and aromatic amino acids in P1 and P1’, which is consistent with aspartyl peptidase activity (Ivry et al., 2017). The majority of the cleavage sites obtained using *Pp*EC15wt were not observed in the double mutant sample, indicating that *Pp*EC15 is an active protease and that mutations in the Asp residues impair its cleavage activity. An iceLogo was used to visualize the specificity profile using the distinct peptide sequences obtained for *Pp*EC15wt (n=156) (Figure 2C). To quantify differences in global substrate specificity between the *Pp*EC15wt and *Pp*EC15dm samples, Z scores from the unique cleavages in each group were also used to generate a difference map derived from residue preferences at each subsite (Figure 2D). This analysis confirmed the preference for Lys at P1 and for aromatic residues at P1’, and it highlighted additional minor preferences at other positions. The profile specificity of *Pp*EC15 includes [L/R]-[K/R/F/Y]-[F/W/Y/R]-[R/M] at positions P2-P1-P1’-P2’, respectively (Figure 2, C-D; Supplemental Figure S1B).

### *Pp*EC15 suppresses plant basal defense responses, and its catalytic activity is not required for all its virulence functions

We were interested in confirming *Pp*EC15’s ability to suppress host defenses and determining if its aspartic protease activity is important for these functions. Callose deposition is a late response to PTI associated with reinforcing cell walls at the site of pathogen attack and is widely used as an indicator of the activation of basal defense responses (Luna et al., 2011). Columbia (Col-0) *Arabidopsis thaliana* (hereafter, Arabidopsis) plants normally exhibit little callose deposition when inoculated with *Pseudomonas syringae* pv. tomato (*Pto*DC3000), whereas extensive callose deposition is observed in leaves inoculated with a disarmed *Pto*DC3000 mutant strain, ΔCEL, which lacks the conserved effector locus (CEL) (Alfano et al., 2000; DebRoy et al., 2004). We therefore transiently expressed *Pp*EC15 in ΔCEL and observed a notable reduction in callose deposition compared to ΔCEL carrying the empty vector (EV) (Figure 3A). We additionally monitored ROS production in the presence of the minimal bacterial flagellin peptide flg22, a known PAMP and elicitor of PTI, in *N. benthamiana* plants transiently expressing *Pp*EC15wt, which demonstrated a significant reduction in ROS production compared to EV controls (Supplemental Figure S2).

**Figure 3.**
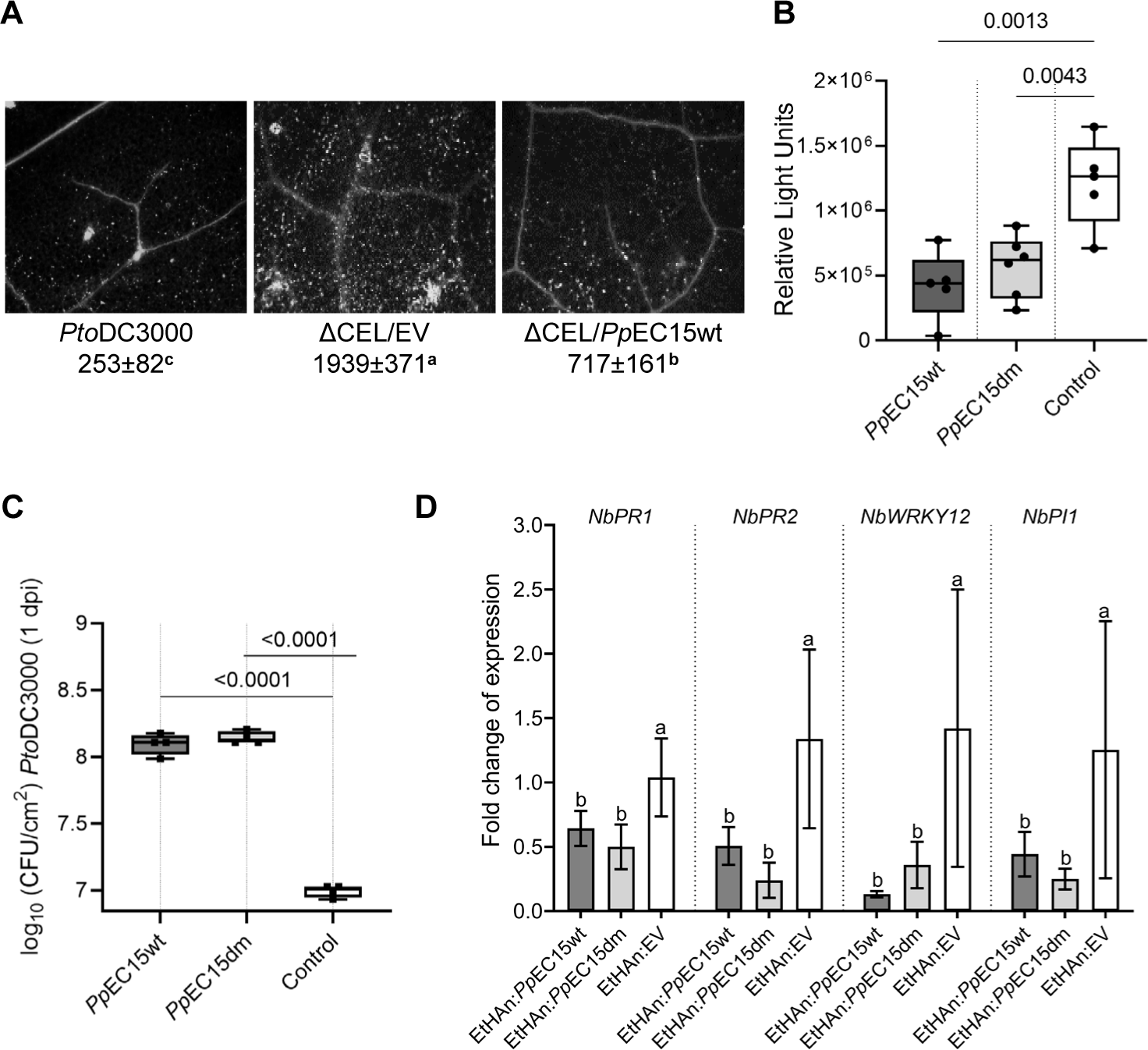
*Pp*EC15 suppresses plant basal defense responses in heterologous plant systems. **A.** Callose deposition in leaves of Arabidopsis Col-0 induced by *Pto*DC3000, ΔCEL/Empty vector (EV) and ΔCEL/*Pp*EC15wt constructs stained with aniline blue. The average number of callose spots ± standard deviation is listed under each representative image. Pairwise *t*-tests were performed and groups with statistically significant differences were denoted with different letters. Representative images are shown (n=24). **B.** Flg22-elicited apoplastic ROS production in *N. benthamiana* transiently expressing *Pp*EC15wt and *Pp*EC15dm 48 hours post-infiltration (hpi). Data are represented as box plots indicating the 25%–75% interquartile range, split by a median line. Whiskers represent maximum and minimum values. Three independent biological replicates were performed. A representative replicate is shown. Statistically significant values (p < 0.05) are determined by a *t*-test using GraphPad Prism 9.0. **C.** *Pto*DC3000 growth in *N. benthamiana* transiently expressing *Pp*EC15wt, *Pp*EC15dm and EV control. Initial inoculum was adjusted uniformly to 10^5^ CFU/mL. Numbers of bacteria were evaluated at 1 dpi. Two independent biological replicates were performed. A representative replicate is shown. Data are represented as box plots indicating the 25%–75% interquartile range, split by a median line. Whiskers represent maximum and minimum values. Statistically significant values (p < 0.05) are determined by a *t*-test using GraphPad Prism 9.0. **D.** Transcript level fold-change of immune marker genes in *N. benthamiana* transiently expressing *Pseudomonas fluorescens* strain EtHan containing EV control, *Pp*EC15wt, or *Pp*EC15dm at 6 hpi. *NbAct1* was used as the internal reference gene. *T*-tests were performed for each comparison and a, b, c were designated groups with statistically significant differences. Two independent biological replicates were performed. Error bars indicate the SD of the technical replicates for each individual sample.

To test whether *Pp*EC15 protease activity was required for immune suppression functions, the double catalytic mutant was included. ROS production and bacterial growth were measured in *N. benthamiana* plants expressing *Pp*EC15wt or *Pp*EC15dm. PAMP-induced ROS production was suppressed in plants transiently expressing the two *Pp*EC15 versions compared to the EV control (Figure 3B). We further inoculated these plants with *Pto*DC3000, which resulted in enhanced bacterial growth when *Pp*EC15wt or *Pp*EC15dm were present (Figure 3C). We additionally measured expression levels of four marker genes associated with SA signaling, *PR1a, PR2*, *WRKY12*, and *PI1* (Noon et al., 2016) in *N. benthamiana* plants inoculated with the non-pathogenic bacterium *Pseudomonas fluorescens* (EtHAn) carrying the EV control, *Pp*EC15wt, or *Pp*EC15dm (Figure 3D). Expression of all marker genes was significantly reduced in the leaves inoculated with EtHAn expressing *Pp*EC15 or *Pp*EC15dm, when compared to leaves inoculated with EtHAn carrying the EV.

To corroborate the results obtained using heterologous systems, we transiently expressed *Pp*EC15wt and *Pp*EC15dm in soybean using *Pto*DC3000 D36E, a polymutant lacking all known Type III secretion effector genes (Figure 4A). Expression of *Pp*EC15wt and *Pp*EC15dm in soybean resulted in enhanced bacterial growth, indicating host immune suppression. To further confirm our results, we generated transgenic soybean lines expressing *Pp*EC15wt under the control of a dexamethasone (DEX)-inducible promoter. Expression of *Pp*EC15 was verified by western blot analysis in two independent transgenic lines 24 hours after induction (Figure 4B). Flg22-induced ROS production was significantly suppressed in the two soybean lines expressing *Pp*EC15 relative to null siblings (Figure 4C). Moreover, the transgenic lines expressing *Pp*EC15 were inoculated with the bacterial pathogen *Pseudomonas syringae* pv. glycinea race 4, which allowed for more bacterial growth compared to the null control (Figure 4D). Together, these data indicate that *Pp*EC15 is a strong suppressor of basal immune responses in Arabidopsis, *N. benthamiana* and soybean, and that its protease activity may not be required for all of its immune suppression functions.

**Figure 4.**
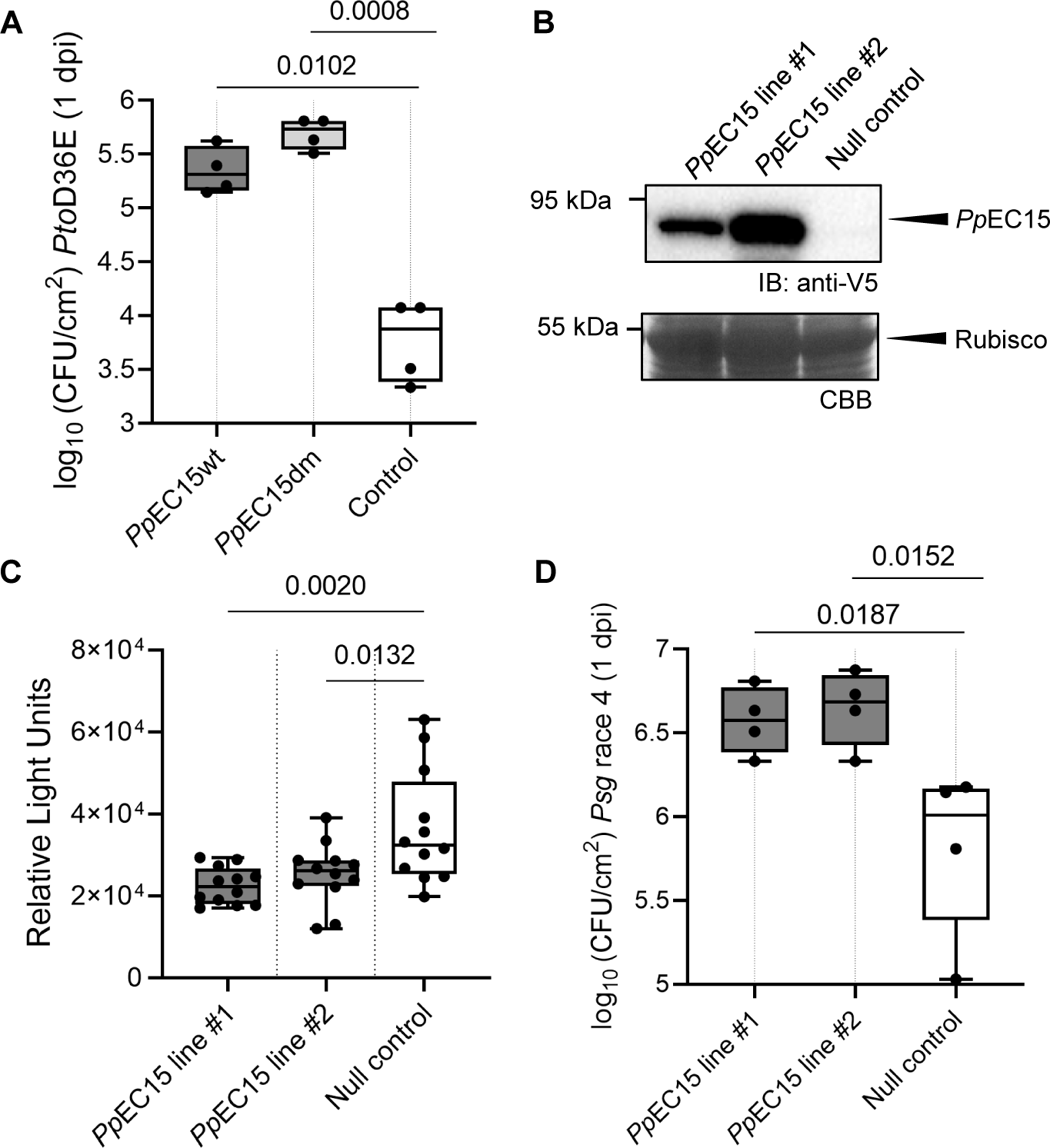
*Pp*EC15 suppresses soybean basal defense responses. **A.** *Pto*DC3000 D36E polymutant lacking all known Type III secretion effector genes growth in soybean transiently expressing *Pp*EC15wt, *Pp*EC15dm and empty vector (EV) control. Initial inoculum was adjusted uniformly to 10^5^ CFU/mL. Numbers of bacteria were evaluated at 1 dpi. Two independent biological replicates were performed with similar results. A representative replicate is shown. Data are represented as box plots indicating the 25%–75% interquartile range, split by a median line. Whiskers represent maximum and minimum values. Statistically significant values (p < 0.05) are determined by a *t*-test using GraphPad Prism 9.0. **B.** Western blot of *Pp*EC15 expression in soybean expression line 1 (*Pp*EC15 line #1) and soybean expression line 2 (*Pp*EC15 line #2) 24 hours after dexamethasone induction. **C.** Flg22-elicited apoplastic ROS production in *Pp*EC15 line #1 and *Pp*EC15 line #2 24 hours after dexamethasone induction. Data are represented as box plots indicating the 25%–75% interquartile range, split by a median line. Whiskers represent maximum and minimum values. Three independent biological replicates were performed. A representative replicate is shown. Statistically significant values (p < 0.05) are determined by a *t*-test using GraphPad Prism 9.0. **D.** *Pseudomonas syringae pv. glycinea* (*Psg*) race 4 growth in transgenic soybean lines expressing *Pp*EC15. Initial inoculum was adjusted uniformly to 10^5^ CFU/mL. Numbers of bacteria were evaluated at 1 dpi. Two independent biological replicates were performed. A representative replicate is shown. Data are represented as box plots indicating the 25%–75% interquartile range, split by a median line. Whiskers represent maximum and minimum values. Statistically significant values (p < 0.05) are determined by a *t*-test using GraphPad Prism 9.0.

### The nuclear localization of *Pp*EC15 is important for its immune suppression functions

To evaluate the function of the predicted NLS, we used fluorescence microscopy to determine the subcellular localization of mVenus-tagged *Pp*EC15 and three mutants (Figure 5A). *Pp*EC15wt-mVenus displayed a strong nuclear and partially cytosolic subcellular localization. In contrast, the *Pp*EC15nls1-mVenus and *Pp*EC15nls2-mVenus mutants both had disrupted nuclear localization, as these proteins became primarily localized to the cytosol (Figure 5A). We additionally tested the subcellular localization of *Pp*EC15dm-mVenus, which showed a similar expression pattern as *Pp*EC15wt-mVenus, suggesting that the catalytic motif substitutions have little influence on the localization of *Pp*EC15 (Figure 5A).

**Figure 5.**
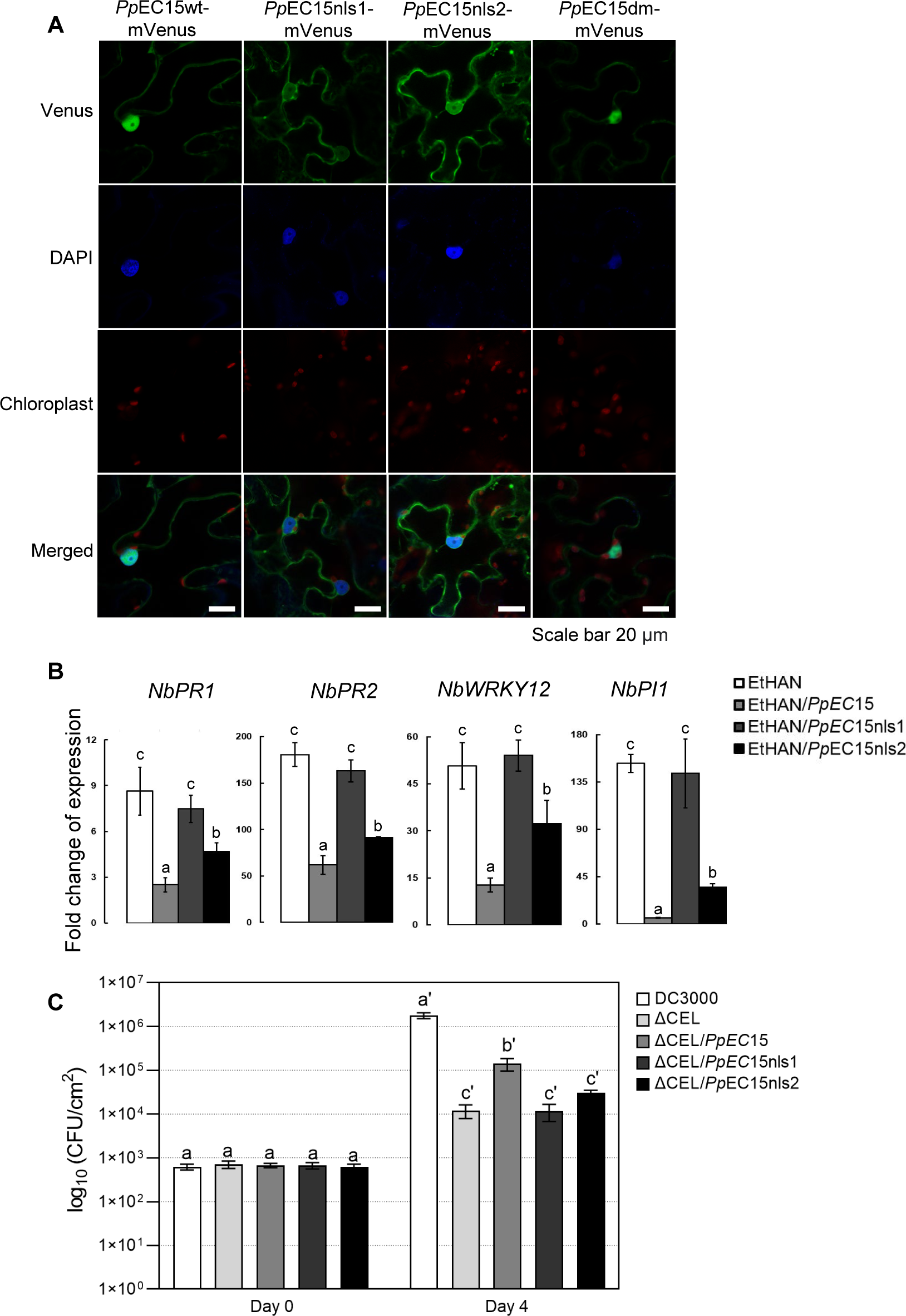
*Pp*EC15 nuclear localization signal is important for its immune suppression functions. **A.** Representative images of the subcellular localization of *Pp*EC15wt, NLS and catalytic mutant variants. Subcellular localization of *Pp*EC15 carboxyl(C)-terminally tagged with Venus was determined by confocal microscopy. All *Pp*EC15-YFP fusions were transiently expressed in *N. benthamiana* by Agrobacterium-mediated infiltration. The leaves were stained with 4′,6-diamidino-2-phenylindole (DAPI) to detect nuclei. Cells were imaged using filter settings that allowed the detection of Venus, DAPI, and chlorophyll autofluorescence. This assay was repeated three times. Photographs were taken at 48 hours post-infiltration (hpi). Scale bars = 20 µm. **B.** Transcript level fold-change of immune marker genes in *N. benthamiana* transiently expressing *Pseudomonas fluorescens* strain EtHan containing empty vector (EV) control, *Pp*EC15wt, or *Pp*EC15 mutant variants at 6 hpi. *NbAct1* was used as the internal reference gene. *T*-tests were performed for each comparison and a, b, c were designated groups with statistically significant differences. Two independent biological replicates were performed. Error bars indicate the SD of the technical replicates for each individual sample. **C.** Bacterial growth *in planta* of *Pto*DC3000, ΔCEL/EV, ΔCEL/*Pp*EC15 or ΔCEL/*Pp*EC15 NLS mutant constructs. Initial inoculum was adjusted uniformly to 10^5^ CFU/mL. Numbers of bacteria were evaluated at 0 dpi and 4 dpi. Two independent biological replicates were performed. Error bars indicate the SD of the technical replicates for each individual sample.

We next aimed to investigate the importance of the NLS for *Pp*EC15 immune suppression functions. We first used an assay in which basal defense triggered by the non-pathogenic bacterium EtHAn suppresses HR induced by subsequent challenge with *Pto*DC3000 (Oh and Collmer, 2005). EtHAn expressing *Pp*EC15 was infiltrated into the *N. benthamiana* leaves (area surrounded by white dashed lines in Supplemental Figure S3). Seven hours later, a partially overlapping area was infiltrated with *Pto*DC3000 (surrounded by red dashed lines in Supplemental Figure S3). The leaf area inoculated with only *Pto*DC3000 developed HR within 36 hours, but the overlapping area did not. In contrast, HR was observed in the overlapping area when EtHAn expressing *Pp*EC15 was used, suggesting that *Pp*EC15 is able to suppress the defense responses elicited by EtHAn (Supplemental Figure S3). Suppression of HR was observed in approximately two-thirds of infiltrated leaf patches expressing *Pp*EC15 compared to control plants which display HR in the entire infiltrated region. *Pp*EC15nls1 and *Pp*EC15nls2 were also expressed from EtHAn and HR suppression was assessed. *Pp*EC15nls1 mutants were unable to suppress basal defenses in this assay, but *Pp*EC15nls2 partially maintained its function with approximately one-third of the leaf area inoculated with EtHAn expressing *Pp*EC15nls2 showed suppressed basal defense.

We next measured expression levels of the defense-related marker genes *PR1a, PR2*, *WRKY12*, and *PI1* (Noon et al., 2016) in *N. benthamiana* plants inoculated with EtHAn carrying EV, *Pp*EC15wt, or *Pp*EC15 NLS mutants (Figure 5B). All marker genes were induced in inoculated leaves, indicating activation of basal defense responses. However, the induction of the four marker genes was significantly lower in the leaves inoculated with EtHAn expressing *Pp*EC15wt or *Pp*EC15nls2, when compared to leaves inoculated with EtHAn carrying the empty vector (EV) or expressing the *Pp*EC15nls1 mutant. These results suggest that *Pp*EC15 can suppress plant defense responses, and this ability is, at least partially, dependent on the NLS.

We additionally tested whether NLS mutations affected *Pp*EC15’s virulence properties in Arabidopsis by quantifying the growth of ΔCEL four days after infection (Figure 5C). ΔCEL carrying *Pp*EC15 grew approximately 10-fold more than ΔCEL carrying EV or *Pp*EC15nls1, while ΔCEL carrying *Pp*EC15nls2 grew approximately 2-fold more than ΔCEL carrying other *Pp*EC15 mutants, suggesting again the *Pp*EC15nls2 mutant partially maintains its suppression functions. Together, our cumulative data indicate that the nuclear localization of *Pp*EC15 is important for its ability to suppress immune activity in *N. benthamiana* and Arabidopsis.

### Yeast-two-hybrid identifies two *Pp*EC15 interacting soybean proteins

To identify soybean proteins that interact with *Pp*EC15, we performed a Y2H screen using a soybean cDNA library. Several candidates were identified, and we validated the interaction of *Pp*EC15 with two of these soybean proteins, PCRF (Peptide Chain Release Factor, Glyma.09G248300) and NAC83 (NAC domain-containing protein 83, Glyma.18g301500) (Figure 6A). Both *Pp*EC15wt and *Pp*EC15dm interacted with PCRF and NAC83, indicating that these interactions are not dependent on functional aspartyl protease activity. Split-luciferase complementation assays further confirmed the interactions between *Pp*EC15wt and *Pp*EC15dm with PCRF (Figure 6B) and NAC83 (Figure 6C). Taken together, our results suggest that mutations in the catalytic motifs weaken *Pp*EC15-soybean protein interactions, but do not abolish it (Figure 6, A-C).

**Figure 6.**
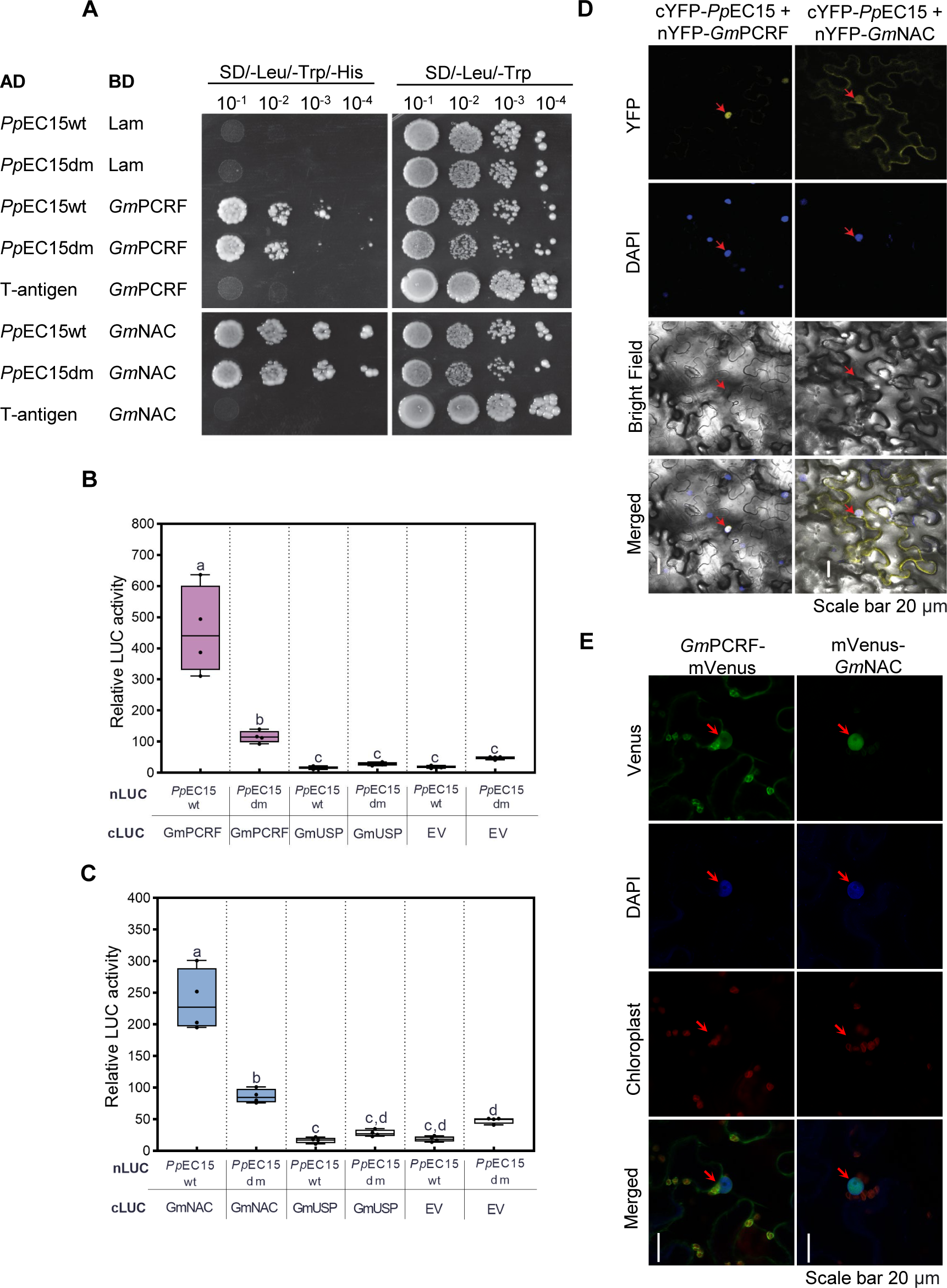
*Pp*EC15 interacts with a soybean transcription factor (*Gm*NAC83) and a soybean peptide chain release factor (*Gm*PCRF). **A.** Yeast-two-hybrid assay showing that *Pp*EC15 interacts with *Gm*PCRF and *Gm*NAC83. The interaction of murine p53 (p53) and SV40 large T-antigen (T) was used as a positive control for the system, and human lamin C (lam) was used as a negative control. **B and C**. Split-luciferase assay of *Pp*EC15 with *Gm*PCRF (B) and *Gm*NAC83 (C). Soybean Universal Stress Protein (USP, Glyma.02G155600) and empty vector (EV) were used as negative control pairs. At 48 h post-Agrobacterium infiltration, the luciferase activity as a result of the association of N and C luciferase fragments through protein-protein interactions were measured. *T*-tests were performed for each comparison and a, b, c were designated groups with statistically significant differences. **D.** Bimolecular fluorescence complementation (BiFC) assay showing that *Pp*EC15 interacts with the soybean proteins *in planta*. DAPI signal was used as a nuclear marker. Arrows indicate nuclei. Representative images are shown (n=20). Scale bars = 20 µm. **E.** Subcellular localization of mVenus-tagged *Pp*EC15 interacting proteins. Green fluorescence indicates the localization of soybean proteins in *N. benthamiana* epidermal cells. Photographs were taken at 48 hours post-infiltration (hpi) using confocal microscopy. Scale bars = 20 µm.

Bimolecular fluorescence complementation (BiFC) analysis was used to confirm the interactions between *Pp*EC15 and the two soybean proteins *in planta* and to investigate where the interactions occur (Figure 6D). Co-expression of *Pp*EC15 with SPL12l (SQUAMOSA promoter-binding-like protein 12 like) (Qi et al. 2016) was used as negative control, which did not result in any fluorescence signal (Supplemental Figure S4). Co-expression of *Pp*EC15 with *Gm*PCRF or *Gm*NAC83 resulted in strong fluorescence in the nuclei (Figure 6D). Strong expression was also observed in the cytoplasm when co-expressing *Pp*EC15 and *Gm*NAC83. When mVenus-tagged *Gm*PCRF was expressed on its own, it localized to both chloroplasts and nuclei (Figure 6E), in line with the predicted chloroplast transit peptide and nuclear localization signal. mVenus-tagged *Gm*NAC83 had strong nuclear localization, consistent with its prediction of a transcription factor (Figure 6E). Together, these results indicate that *Pp*EC15 interacts with two of its host targets in the nucleus and cytoplasm, consistent with its subcellular localization.

### Proximity labeling identifies novel *Pp*EC15 interacting proteins

To further investigate interactions between *Pp*EC15 and soybean proteins, we performed a proximity labeling (PL) assay by fusing *Pp*EC15wt, *Pp*EC15dm, and an NLS-GFP control to the miniTurbo biotin ligase (Figure 7A) (Branon et al. 2018; Zhang et al. 2020; Zhang et al. 2019). The fusion proteins were transiently expressed in *N. benthamiana*. Capture of the biotinylated proteins (Supplemental Figure S5) enabled the identification of the enriched interacting proteins by LC–MS/MS (Supplemental Table S1). Importantly, *Pp*EC15 peptides were detected in the dataset, confirming its expression *in planta* (Supplemental Table S1). We identified 334 proteins exclusively enriched in the *Pp*EC15wt dataset compared to *Pp*EC15dm, and 44 proteins enriched both in the wild-type and mutated versions compared to the GFP control (Figure 7B), suggesting that mutation in the catalytic motifs impairs *Pp*EC15 ability to interact with most of its partners, but it does not completely abolish it. This is consistent with the Y2H screens, protease specificity and split-luciferase complementation assays. PL identified several proteins previously shown to be involved in biotic stress responses in other pathosystems, including a calmodulin-binding protein (Niben101Scf06222g01010.1), a WRKY transcription factor (Niben101Scf03786g04014.1), and a topless-related protein (Niben101Scf05720g06001.1) (Figure 7C). The classification of the potential *Pp*EC15 interactors into putative functional categories revealed enrichment in biological processes related to chloroplast function and transcription (Figure 7D), which is consistent with the localization of the previously identified interacting partners of *Pp*EC15.

**Figure 7.**
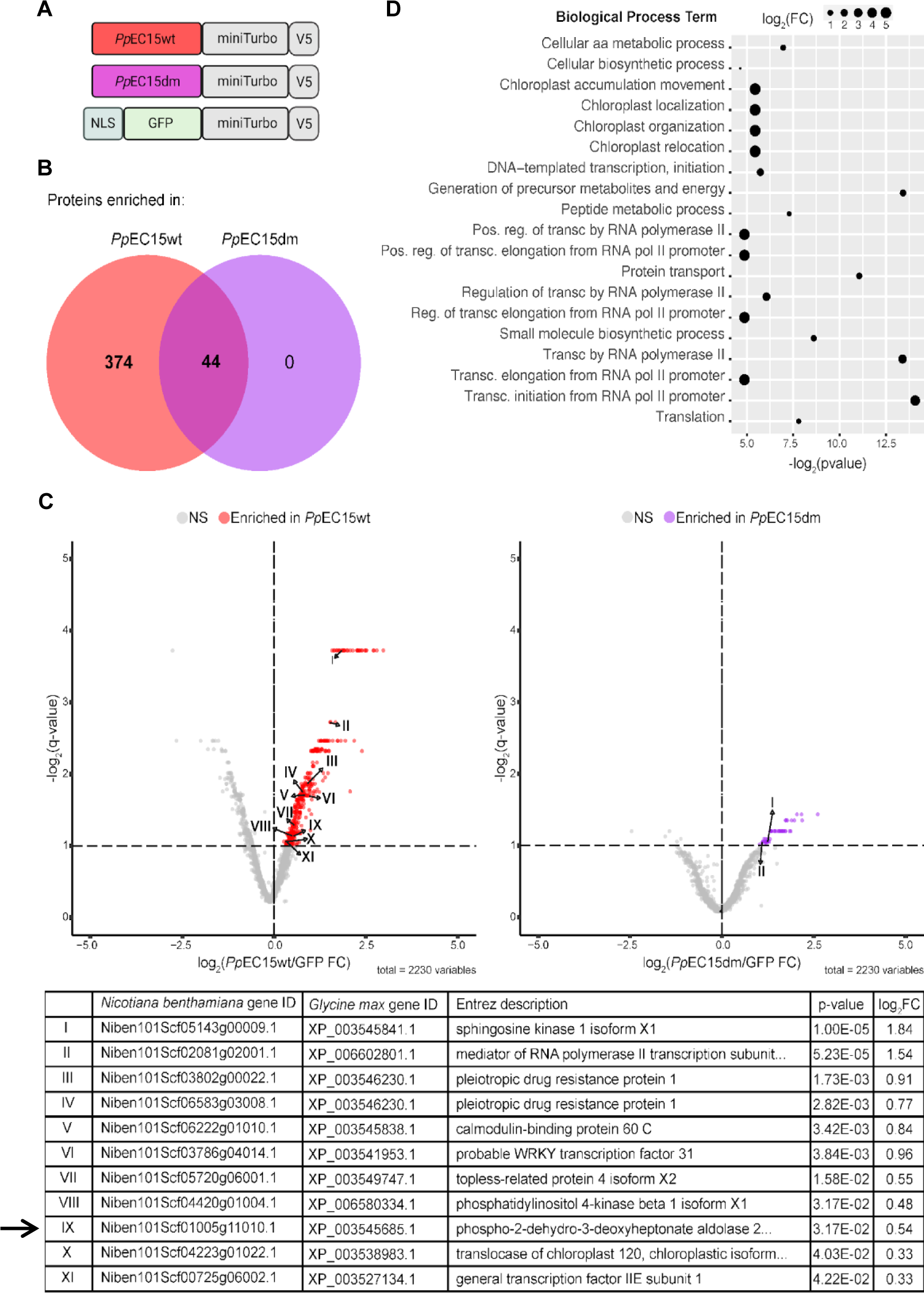
Proximity-based labeling assay identifying novel *Pp*EC15 potential interacting partners. **A.** Schematic representation of the constructs used for the transient expression in *N. benthamiana*. *Pp*EC15wt, *Pp*EC15dm, and a nuclear-localized GFP control were fused to the amino terminus of the miniTurbo biotin ligase. **B.** Venn diagram of the number of proteins enriched in the *Pp*EC15wt and *Pp*EC15dm dataset compared to the GFP control. **C.** Volcano plot of *Pp*EC15wt and *Pp*EC15dm proximity labeling analysis highlighting a selection of possible direct targets. **D.** Selected biological processes enriched among *Pp*EC15wt and *Pp*EC15dm interacting proteins.

Interestingly, one of the proteins enriched in this assay, a 2-dehydro-3-deoxyphosphoheptonate aldolase (Niben101Scf01005g11010.1), is involved in the biosynthesis of SA and secondary metabolites, and it has a homologous protein in soybean with 70.6% amino acid sequence similarity (*Gm*DAHP, Glyma.14g176600.1). *Gm*DAHP was previously identified in our initial Y2H screen, but the direct interaction was not confirmed. The role of SA in plant defense against biotic stresses is well established (Lefevere et al., 2020). Therefore, we sought to confirm the interaction between *Pp*EC15 and *Gm*DAHP. BiFC and split-luciferase complementation assays confirmed that *Pp*EC15 and *Gm*DAHP interact, and that the interaction occurs in the nucleus and the chloroplasts (Figure 8, A-B). *Gm*DAHP has a strong predicted chloroplast transfer peptide and appears to associate with chloroplasts when expressed alone (Figure 8C). Since *Pp*EC15 displayed strictly nuclear-cytoplasmic localization when expressed on its own, we were curious if the interaction in the chloroplast was an artifact of the BiFC assay, which involves an irreversible reconstitution of YFP. To investigate this, we performed a co-expression assay with *Pp*EC15-RFP and *Gm*DAHP-GFP. Our results show that although *Gm*DAHP still localizes in chloroplasts in the presence of *Pp*EC15, it appears that when they are co-expressed, *Gm*DAHP forms small aggregates in the cells (Supplemental Figure S6), suggesting that *Pp*EC15 could affect *Gm*DAHP stability or intracellular dynamics. Interestingly, in the presence of DAHP, *Pp*EC15 also localized to some chloroplasts. It is possible that *Pp*EC15 is imported to chloroplasts in a “piggyback” manner, suggesting that distinct mechanisms of host immune suppression might be explored by *Pp*EC15.

**Figure 8.**
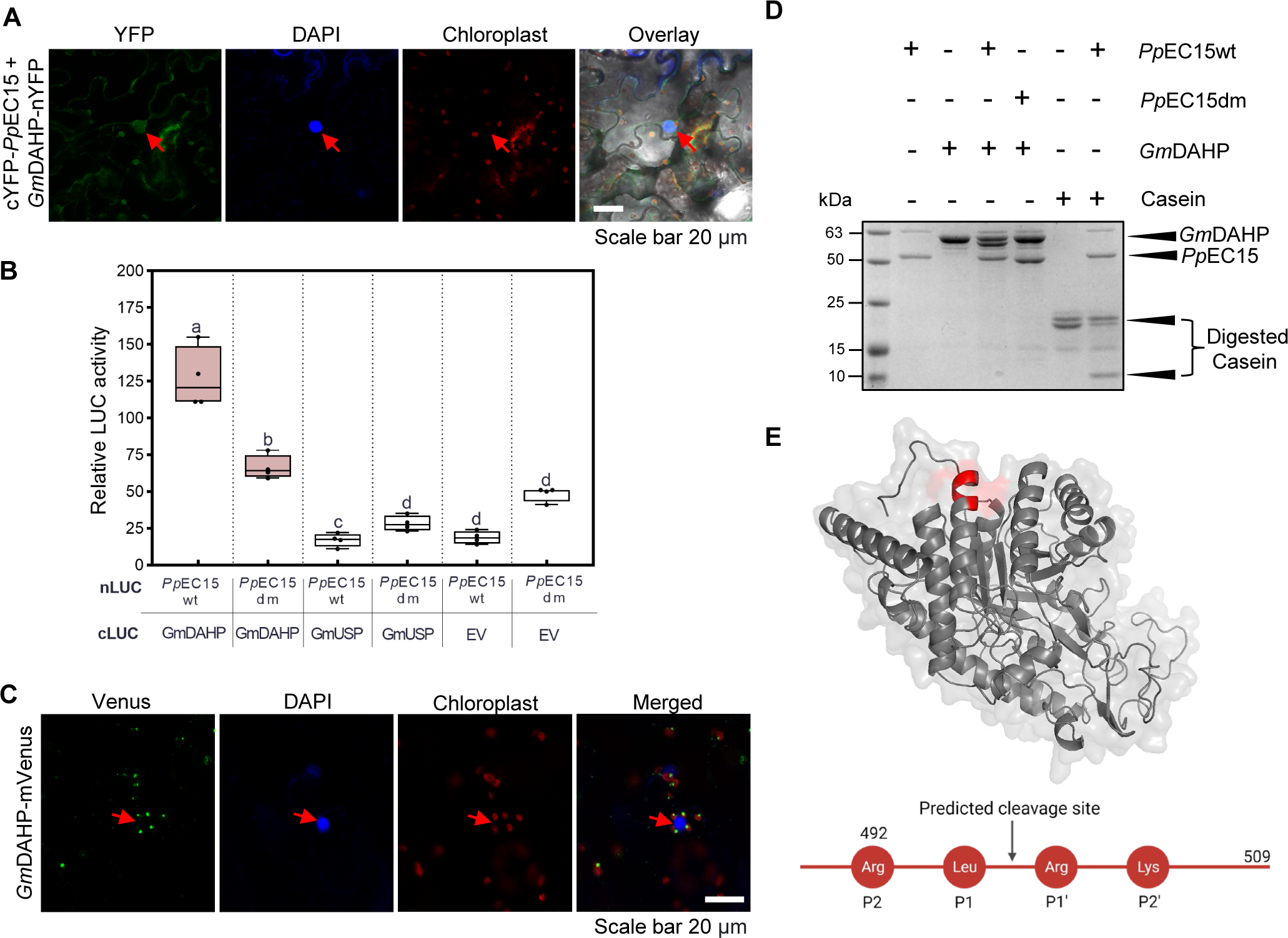
*Pp*EC15 interacts with and cleaves a soybean Class-II DAHP synthase (*Gm*DAHP). **A.** BiFC assay showing that *Pp*EC15 interacts with *Gm*DAHP *in planta*. DAPI signal was used as a nuclear marker. Arrows indicate nuclei. Representative images are shown (n=20). Experiments were conducted three times with the same results. **B.** Split-luciferase assay of *Pp*EC15 with *Gm*DAHP. Soybean Universal Stress Protein (USP, Glyma.02G155600) and empty vector (EV) were used as negative control pairs. At 48 h post-Agrobacterium infiltration, the luciferase activity as a result of the association of N and C luciferase fragments through protein-protein interactions were measured. *T*-tests were performed for each comparison and a, b, c were designated groups with statistically significant differences. **C.** Subcellular localization of mVenus-tagged *Gm*DAHP. Green fluorescence indicates the localization of the soybean protein in *N. benthamiana* epidermal cells. DAPI signal was used as a nuclear marker. Arrows indicate nuclei. Representative images are shown (n=20). Photographs were taken at 48 hours post-infiltration (hpi) using confocal microscopy. **D.** SDS-PAGE demonstrating the cleavage assay performed with *Pp*EC15wt and *Pp*EC15dm with *Gm*DAHP. 1 µg of recombinant *Pp*EC15 protein and 2 µg of the recombinant *Gm*DAHP protein were incubated at 37 °C for 16 hours. Proteins were visualized using Coomassie brilliant blue (CBB) staining. Casein was used as a model substrate. **E.** DAHP structural modeling using the Phyre2.0 algorithm using *Pseudomonas aeruginosa* phospho-2-dehydro-3-deoxyheptonate aldolase; (PDB, 1dpj) as a template with 100% confidence score and 62% coverage. The predicted cleavage site is identified in red.

### *Pp*EC15 cleaves *Gm*DAHP

To further investigate how *Pp*EC15 manipulates these soybean proteins, we performed an *in vitro* cleavage assay using *Pp*EC15wt, *Pp*EC15dm and the three soybean interactors, *Gm*NAC83, *Gm*PCRF, and *Gm*DAHP. Recombinantly-produced *Pp*EC15wt and *Pp*EC15dm were incubated with the interactors and cleavage was assessed by SDS-PAGE after 16 h of incubation (Supplemental Figure S7; Fig 8D). No digestion products were observed for NAC83 or PCRF (Supplemental Figure S7), indicating the *Pp*EC15 does not cleave these targets. Strikingly, cleavage of DAHP by *Pp*EC15 was detected based on the presence of a ∼54 kDa peptide, suggesting that the cleavage occurs near the amino (N) or carboxyl (C)-terminus (Figure 8D). Consistent with the predicted cleavage site motif we identified (Figure 2, D-E; Supplemental Table S2), *Gm*DAHP contains Arg-Leu-Arg-Lys at positions P2, P1, P1’, P2’ (amino acids 492-495), respectively (Figure 8E). We hypothesize that this is the position where the cleavage of *Gm*DAHP occurs, such that the larger fragment generated would be ∼54 KDa compared to full length ∼58 KDa protein (Figure 8D), although this remains to be experimentally validated.

### Soybean-interacting proteins are involved in plant defense responses

To determine if the *Pp*EC15-interacting soybean proteins are involved in plant immunity, we generated bean pod mottle virus (BPMV) constructs to silence their expression (Figure 9, A-B). We assayed changes in the dynamics of flg22-elicited ROS production and SA production. Silencing of *GmPCRF*, *GmNAC83* and *GmDAHP* led to reduced flg22-induced ROS production, suggesting that these genes might act as positive immune regulators (Figure 9C). Furthermore, silencing of *GmDAHP* led to a significant reduction in the production of SA, again suggesting a possible immune function (Figure 9D). These results suggest that *Gm*PCRF, *Gm*DAHP, and *Gm*NAC83 might play a role in plant defenses against SBR.

**Figure 9.**
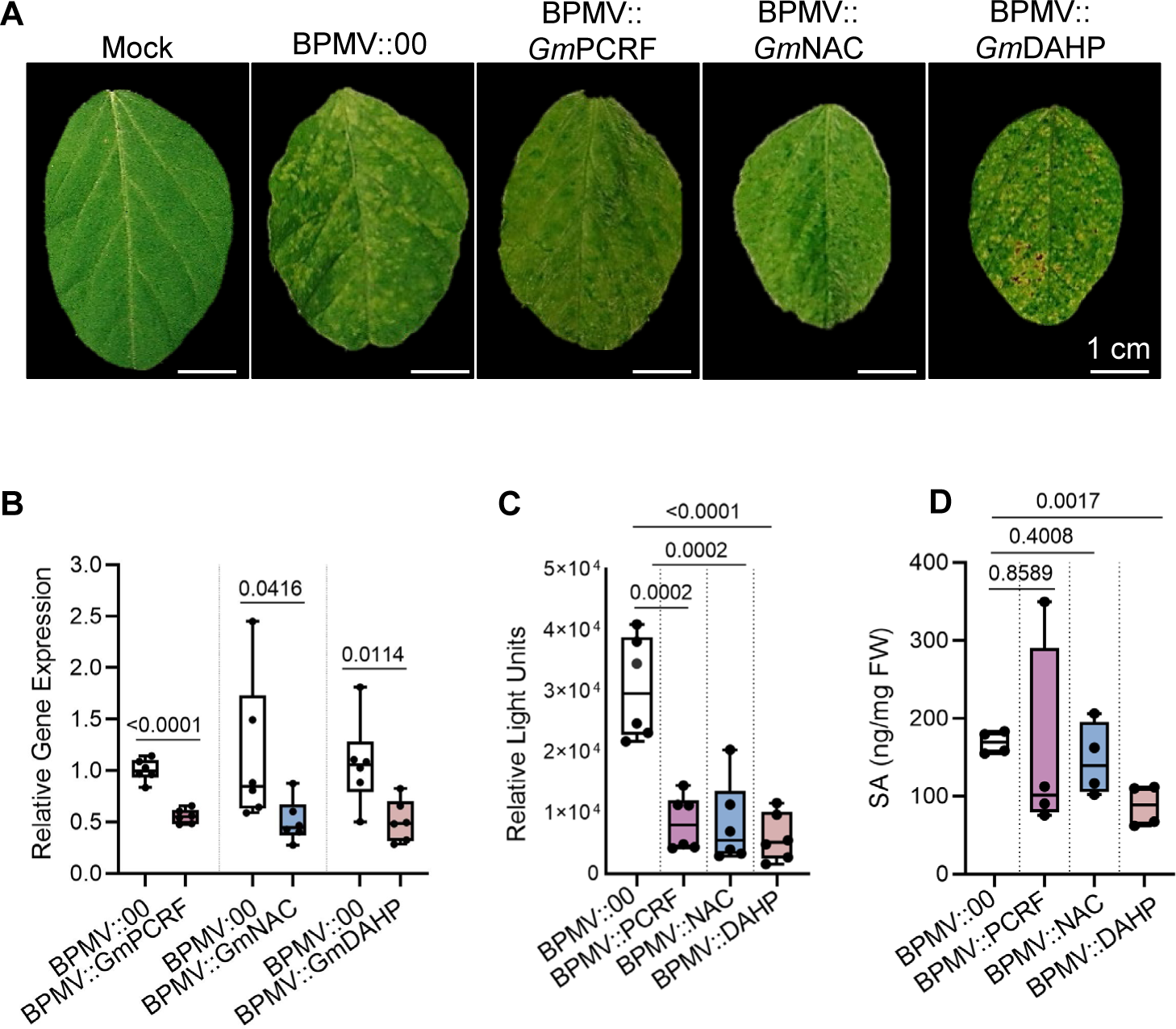
Silencing of *Pp*EC15 interactors alters flg22-induced apoplastic ROS production and salicylic acid (SA) production. **A.** Virus-induced gene silencing (VIGS) mediated by bean pod mottle virus (BPMV) was used to knock down the expression of *Gm*PCRF (Peptide Chain Release Factor, Glyma.09G248300), *Gm*NAC83 (NAC domain-containing protein 83, Glyma.18g301500), and *Gm*DAHP (Class-II DAHP synthase, Glyma.14g176600) in soybean. Representative pictures of BPMV-infected soybean and control at 21 dpi are shown. **B.** RT-qPCR demonstrating the silencing of the soybean interactors 14 days after viral infection. *GmNREG3* (Glyma.08G211200) was used as the internal reference gene. Data are represented as box plots indicating the 25%–75% interquartile range, split by a median line. Whiskers represent maximum and minimum values. Statistically significant values (p < 0.05) are determined by a *t-*test using GraphPad Prism 9.0. The experiment was conducted at least three times with similar results. **C.** Flg22-elicited apoplastic ROS production in BPMV::00 (empty vector), BPMV::PCRF, BPMV::NAC83, and BPMV::DAHP-infected plants 14 days after viral infection. Data are represented as box plots indicating the 25%–75% interquartile range, split by a median line. Whiskers represent maximum and minimum values. A *t-*test was performed for the pairwise comparisons with the control group BPMV:00. Significant differences based on Student’s *t*-test are indicated by the p-values shown in the graphs. Three independent experiments with two technical replicates were performed with similar results. A representative replicate is shown. **D.** SA production in BPMV::00, BPMV::PCRF, BPMV::NAC83 and BPMV::DAHP-infected plants 14 days after viral infection. Data are represented as box plots indicating the 25%–75% interquartile range, split by a median line. Whiskers represent maximum and minimum values. Statistically significant values (p < 0.05) are determined by a *t*-test for the pairwise comparisons with the control group BPMV:00. Two independent biological replicates were performed. A representative replicate is shown.

## Discussion

In this study, we examined the *in vitro* and immune suppressing properties of *Pp*EC15, which was previously identified in a screen for haustoria-expressed, secreted proteins that suppressed immunity in heterologous expression systems (Qi et al., 2018). Prior motif analysis predicted a secretion signal peptide and NLS in the N-terminus of *Pp*EC15, and an aspartyl protease family A1A domain (Link et al., 2014; Qi et al., 2018). Our work suggests that the NLS is important for immune suppression functions, but the protease activity was not required in the immune suppression assays we performed.

The identification and characterization of conserved effectors with functions in suppressing or eliciting the plant immune system and the identification of their immune targets is critical to generating durable resistance against SBR. In a recent study, a *P. pachyrhizi* effector, Phapa-7431740, suppressed PTI and interacted with a soybean glucan endo-1,3-β-glucosidase (*Gm*βGLU), a PR protein belonging to the PR-2 family (Bueno et al., 2022). Similarly, *Pp*EC23 was previously shown to suppress soybean PTI and ETI and physically interact with the soybean transcription factor SQUAMOSA-PROMOTER BINDING PROTEIN-LIKE (SPL)12l, a negative immune regulator (Qi et al., 2016) and possible susceptibility gene. Despite the importance of this pathogen to soybean production, the characterization of *P. pachyrhizi* effectors is limited to the two above-mentioned studies.

We used heterologous plant systems and the host species soybean to demonstrate that *Pp*EC15 is a strong suppressor of basal defense responses elicited by a non-pathogenic bacterium, callose deposition, ROS production, and expression of defense-related genes, enhancing susceptibility to bacterial pathogens (Fig 3, A-D; Figure 4, A-D; Supplemental Figure S2-3). *P. pachyrhizi* effectors may target components of PTI, ETI, or both, as there is increasing evidence of a complex interplay between PRR-mediated and NLR-mediated signaling (Yuan et al., 2021). Because *Pp*EC15 affected the expression of defense marker genes *PR1a*, *PR2*, *WRKY12*, and *PI1* in response to bacterial infection (Figure 3D; Figure 5B), we expect that it affects regulatory components that activate expression of these genes. Moreover, the high level of conservation across *P. pachyrhizi* isolates and other rust species, including economically important pathogens of wheat, oat and poplar, suggests that *Pp*EC15 may have conserved functions in *P. pachyrhizi* virulence and likely in other rust fungi.

As an initial step to understanding how *Pp*EC15 disrupts host cell function, we investigated its subcellular localization. In agreement with the NLS prediction, *Pp*EC15-mVenus was present in nuclei, and it was also detected in the cytosol (Figure 5A). *Pp*EC15-interacting proteins, *Gm*PCRF, *Gm*NAC83, and *Gm*DAHP were localized to different compartments in *N. benthamiana* epidermal cells: *Gm*PCRF (nuclear and cytoplasmic), *Gm*NAC83 (nuclear), and *Gm*DAHP (chloroplast) (Figure 5C; Figure 7B). When co-expressed with *Pp*EC15 in BiFC, the interactions were mainly observed in the nuclei, but fluorescence was also detected in chloroplasts and cytosol, although to a lesser extent (Figure 6D; Figure 8A). While these results suggest that interactions with *Pp*EC15 can cause relocalization of host proteins, interactions in BiFC are virtually irreversible (Kodama and Hu 2012), and the relocalization could be an artifact of the assay such that once the association occurs, *Pp*EC15 pulls its targets to the nucleus where it normally localizes. Although our subcellular localization results show that *Pp*EC15 is not able to relocalize DAHP, it appears that *Pp*EC15 induces DAHP protein agglomeration near to the nucleus (Supplemental Figure S6). Remarkably, our work shows that *Pp*EC15 can cleave *Gm*DAHP, which could be causing the agglomerates observed when the two are co-expressed. DAHP is the first enzyme in the shikimate pathway, which plays a central role in the biosynthesis of defense metabolites in plants (Maeda and Dudareva, 2012). It remains to be determined if cleavage of DAHP would cause the enzyme to become inactive or affect its immune functions.

Arogenate dehydratase 6, an enzyme involved in the shikimate pathway, is differentially expressed upon SBR infection in susceptible and resistant soybean lines (Hossain et al., 2018). Furthermore, plant genes encoding enzymes in the shikimate pathway are differentially regulated in response to several environmental cues, including wounding and pathogens (Görlach et al., 1995; Zhao and Last, 1996). DAHP synthase genes are differentially expressed during pathogen attack in Arabidopsis (Keith et al., 1991), *Solanum lycopersicum* (tomato) (Görlach et al., 1995), and *Solanum tuberosum* (potato) (Jones et al., 1995). Similarly, expression of *Gh*DAHP in cotton is induced upon infection with *Verticillium dahliae*, and overexpression of *Gh*DAHP in Arabidopsis transgenic plants led to resistance against Verticillium (Yang et al., 2015). Pathogen effectors can target components of phytohormone signaling pathways to circumvent the host immune system. For example, *Ustilago maydis, Verticillium dahliae,* and *Phytophthora sojae* attenuate plant SA biosynthesis by degrading SA precursors chorismate or isochorismate using the effectors Cmu1, VdIsc1, and PsIsc1, respectively (Djamei et al., 2011; Liu et al., 2014). Furthermore, silencing of *GmDAHP* led to suppression of flg22-induced apoplastic ROS production and a significant reduction in SA production in soybean (Figure 9, C-D). Our work suggests that *Pp*EC15 may target the shikimate pathway to contribute to *P. pachyrhizi* virulence, in part, by perturbing the activity of *Gm*DAHP likely mediated by its cleavage. DAHP is highly conserved across various organisms (Tzin et al. 2012), and has been shown to have a dimer-of-dimers homotetrameric structure (Figure 10A) (Balachandran et al. 2016). Mutation analysis of *E. coli* DAHP synthase shows that the termini of the enzyme play a critical role in the formation of the dimeric structure required for its functionality (Xu et al. 2004). Because DAHP forms tetramers, cleavage of the termini could potentially prevent the formation of the dimeric complex, impairing its functionality and subsequent production of SA. Although there are no known proteases that recognize cleavage sites in protein helices, a surprising amount of protease cleavage sites have been mapped in helices on folded proteins (Barkan et al. 2010). This suggests that proteins undergo conformational changes and that the helices are folding and unfolding enough to adopt a structure that the protease can recognize. Moreover, the helices are more prone to unfolding near the termini of the protein.

**Figure 10.**
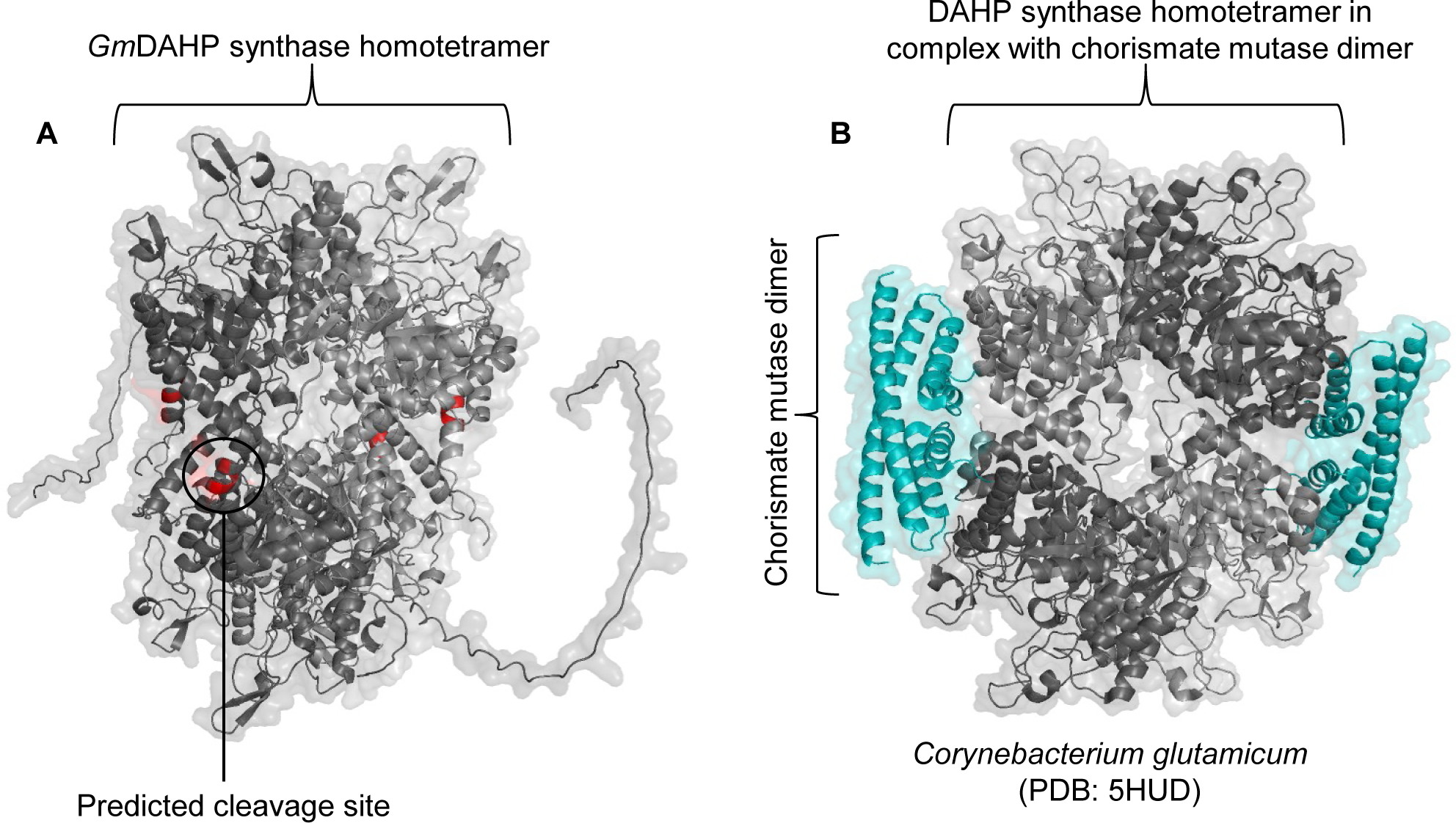
**A.** DAHP homotetramer structural modeling predicted through AlphaFold multimer and visualized using PyMOL Molecular Graphics System. The predicted cleavage site is identified in red. **B.** Homotetramer of DAHP from *Corynebacterium glutamicum* in complex with a dimer of chorismate mutase identified in blue cyan (PDB: 5HUD).

Furthermore, the homotetramer DAHP was shown to interact and form a complex with a dimer of chorismate mutase, a key enzyme in the shikimate pathway that displays low activity without its interaction with DAHP (Sasso et al. 2009). Interestingly, the predicted *Pp*EC15 recognition cleavage site is located at an interaction interface of the DAHP-chorismate mutase complex (Figure 10B). It is possible that cleavage of the C-terminus of DAHP could affect the formation of the DAHP-chorismate mutase complex, leading to a reduced activity of chorismate mutase, therefore reducing downstream SA production. Subsequent studies aiming at elucidating the functionality of the truncated DAHP form, as well as its potential role in mediating plant defense responses will be of significant interest. It is noteworthy that chloroplastic proteins were enriched in our PL experiment. Although *Pp*EC15 is nucleo-cytoplasmic, *Pp*EC15 and DAHP interact in chloroplasts and in nuclei in BiFC and our co-localization assay showed that *Pp*EC15 partially localizes to chloroplasts in the presence of DAHP. It is possible that, despite lacking an identifiable chloroplast transit peptide, *Pp*EC15 can move to these organelles in a “piggyback” fashion to manipulate chloroplastic proteins to subvert plant immunity.

Interestingly, despite being an effector protease, our protein-protein interaction assays using the inactive *Pp*EC15 (*Pp*EC15dm) harboring mutations in the two catalytic Asp residues, show that *Pp*EC15dm still maintains the interaction with the soybean partners to a certain extent, suggesting that the catalytic activity is not required for the interaction. Oftentimes, mutations on catalytic residues of proteases stabilize association with their substrates, presumably because they are unable to cleave substrates and release them. Additionally, as the catalytically inactive mutant was still capable of suppressing basal immune responses including flg22-elicited ROS and defense-related marker genes and led to increased bacterial growth in *N. benthamiana* and in soybean, this activity does not appear to be important for *Pp*EC15’s activity in this context.

*Gm*NAC83 and *Gm*PCRF were identified as *Pp*EC15 interacting partners in our Y2H screen and confirmed using BiFC and split luciferase assays in *N. benthamiana*. NAC proteins are a plant-specific transcription factor family well-known for their roles as regulators of plant defense responses (Puranik et al., 2012). In a transcriptome analysis of susceptible BRS184 soybean and *Rpp3* (*Resistance to Phakopsora pachyrhizi 3*) near-isogenic lines, NAC transcription factors were highly expressed and upregulated during incompatible interactions (Hossain et al., 2018). Moreover, in Arabidopsis, PCRF is required for the proper translation, stability, and normal processing of UGA-containing polycistronic transcripts in chloroplasts (Meurer et al., 2002). In cucumber, both SA treatment and pathogen inoculation induced the expression of a putative translation release factor, suggesting a potential role for the gene products in systemic acquired resistance or pathogen resistance (Bovie et al., 2004). Here, we demonstrate that silencing of *GmNAC83* or *GmPCRF* leads to reduced flg22-induced apoplastic ROS production (Figure 9B), suggesting that these two proteins might positively regulate soybean immunity.

A number of the novel putative interactors identified in our PL assay have been previously implicated in biotic stress responses in multiple pathosystems and include a calmodulin-binding protein (Niben101Scf06222g01010.1), a WRKY transcription factor (Niben101Scf03786g04014.1) and a topless-related protein (Niben101Scf05720g06001.1) (Wani et al., 2021; Yuan et al., 2021; Saini and Nandi, 2022). WRKY transcription factors have been shown to play a role in soybean-*P. pachyrhizi* interactions (Pandey et al., 2011; Bencke-Malato et al., 2014). Transcriptomic analysis of resistant and susceptible SBR-infected soybean plants revealed that WRKY transcription factors were overrepresented during infection, and their expression profiles were consistent with defense and secondary metabolism genes (van de Mortel et al., 2007). It will be interesting to confirm the direct interaction between these novel putative candidates and *Pp*EC15 and to determine if the effector is modulating these immune-related proteins through proteolytic cleavage. Notably, neither *Gm*NAC83 or *Gm*PCRF were identified in our PL assay for several possible reasons, including an inappropriate time-point to capture the interaction, low protein abundance, low-efficiency labeling, lack of surface-exposed lysines, or the lack of a specific stimulus that would trigger the interaction considering that a heterologous plant system was used.

Interestingly, approximately 20% (70) of the proteins enriched in our miniTurbo dataset using *Pp*EC15wt contained one of the potential combinations of the preferential cleavage sites between a basic (P1) and a hydrophobic (P1’) amino acid [L/R][K/R/F/Y][F/W/Y/R][R/M] (Figure 2, D-E; Supplemental Table S2). For example, *Nb*DAHP harbors a predicted *Pp*EC15 cleavage motif (RKRR) at its C-terminus which is also present in the soybean homolog cleaved by *Pp*EC15 (RKRL). While *Pp*EC15 can cleave one of its interactors, our data indicate that this proteolytic activity is not required for at least some of the effector virulence functions. We hypothesize that *Pp*EC15 may be a multifunctional effector protein with aspartyl protease activity that also manipulates other host proteins so they can no longer serve their functions in immune signaling. The mechanism by which these interactions occur remains to be explored.

The identification and characterization of effector proteins that might be essential or play important roles in pathogen virulence are of crucial importance in the development of strategies to breed or engineer durable resistance against SBR. The conservation of *Pp*EC15, its ability to suppress host defense responses and the potential to identify a consensus cleavage site indicate that it may be a good target in the use of new emerging technologies to engineer disease resistance such as the engineering of pathogen protease decoys that can lead to activation of soybean resistance genes upon effector protease target cleavage (Kim et al., 2016; Helm et al., 2019; Pottinger et al., 2020). Furthermore, strategies to generate disease resistance based on the targeting of conserved effectors is more likely to be durable (Dangl et al., 2013), which requires a better understanding of effectors secreted by plant pathogens and their interactions with host plant proteins.

## Materials and Methods

A detailed description of materials and methods used in this manuscript are provided in

Supplemental File 1.

### Sequence alignment and protein modeling

*Pp*EC15 full-length sequence was used as a query in NCBI BLAST (https://www.ncbi.nlm.nih.gov/). Sequences from *Pp*EC15 homologs were then used in MEGA7 software (Kumar et al., 2016) for multiple sequence alignment. The crystal structure of *Pp*EC15 and DAHP were modeled using the PyMOL Molecular Graphics System.

### Construction of *Pp*EC15 mutants

Overlapping PCR, using primers that harbor desired mutations in the NLS sequence or catalytic motifs, were used to generate the *Pp*EC15 mutants (primers listed in Supplemental Table S3). The full-length mutagenized DNA fragments were amplified using full-length primers and were subsequently cloned into pCR™8/GW/TOPO™ as the entry clones, sequence verified, and used for further applications.

### *Pp*EC15-casein cleavage assay

pGEX-4T1:GST-*Pp*EC15 and pGEX-4T1:GST-*Pp*EC15dm fusion proteins were expressed in *Escherichia coli* Lemo21 (DE3) and purified. To eliminate the GST-tag effect on the protease activity, GST was removed with the Thrombin CleanCleave^TM^ Kit (Sigma-Aldrich) following the manufacturer instructions. The casein digestion assay was performed with the purified native *Pp*EC15 proteins, wild type (wt) and double mutant (dm) using 2 µg of casein with 1 µg of *Pp*EC15 variants in PBS buffer (140 mM NaCl, 2.7 mM KCl, 10 mM Na_2_HPO_4_, 1.8 mM KH_2_PO_4_, pH 7.3) incubated at 37 °C for 16 hours. Proteins were visualized using Coomassie brilliant blue (CBB) staining.

### Growth of bacterial strains and triparental matings

*E. coli* and *Agrobacterium tumefaciens* were grown in Luria-Bertani (LB) broth at 37°C or 28°C, respectively. *Pseudomonas* spp. strains were grown in either LB or King’s B (KB) medium at 28 °C. When appropriate, antibiotics were included at the following concentrations: rifampicin (25 µg/mL), kanamycin (50 µg/mL), ampicillin (100 µg/mL, gentamicin (25 µg/mL). Plasmids were mobilized from *E. coli* to *Pseudomonas* strains by triparental mating using *E. coli* HB 101 (pRK2013) as a helper strain.

### Plant material

*N. benthamiana* and soybean (*Glycine max* cv. Williams 82) plants were grown in controlled environment chambers at an average temperature of 24 °C (range 20 °C-26 °C), with 45-65% relative humidity under long-day conditions (16 h light). Arabidopsis plants were grown in controlled environment chambers at an average temperature of 22 °C (range 18 °C-24 °C), with 45%-65% relative humidity under short-day conditions (10 h light). All plants were grown in LC-1 potting soil mix (Sungro horticulture, Catalog Number: 504307), and were fertilized once a week with 15-5-15 Cal-Mag Special (Peters Excel, G99140).

### Bacterial inoculations and *in planta* growth

Four to six-week-old *N. benthamiana* or 4–5-week-old Arabidopsis were infiltrated with bacterial suspensions using a needleless syringe. *A. tumefaciens* strains were resuspended in an infiltration buffer (10 mM MES (pH 5.6), 10 mM MgCl_2_, 100 µM acetosyringone) and kept at room temperature for 3 h before infiltration. *P. fluorescens* strain EtHAn expressing *Pp*EC15 variants were inoculated at 2 × 10^8^ CFU·ml^-1^ and *P. syringae* strain DC3000 at 2 × 10^7^ CFU·mL^-1^ in 10 mM MgCl_2_ prior to infiltration. Bacterial proliferation was determined by cutting leaf disks with a cork borer (inner diameter 0.5 cm) and homogenizing in 50 μl of inoculation buffer prior to plating on KB plates containing appropriate antibiotics.

### Callose staining and microscopic analysis

Arabidopsis Col-0 leaves were harvested 12 h after bacterial infiltration, cleared, and stained with aniline blue for callose as previously described (Adam and Somerville, 1996). Leaves were evaluated with a Zeiss Axioplan II microscope with an A4 fluorescence cube. Quantification of callose depositions was determined using ImageJ software (NIH). Six adjacent fields along the leaf length (avoiding the midvein, leaf edge, or syringe-damaged area) were analyzed and averaged.

### ROS burst assay

ROS burst assays were adapted from a previously established protocol for Arabidopsis (Bredow et al., 2019). Briefly, leaf discs were collected from soybean plants 14 days after BPMV inoculation or from transgenic soybean plants 24 hours after DEX-induction in 96-well plates containing 100 µL of water. The next day, water was replaced with 100 µL of elicitor solution containing 100 µM of the luminol (Sigma Aldrich), 10 µg/mL of horseradish peroxidase (HRP) (Sigma Aldrich), and 100 nM of flg22 (VWR). Luminescence measurements were taken on a Promega GloMax plate reader every 2 min for 40 min with 1000 ms integration time.

### Generation of stable transgenic soybean lines expressing a dexamethasone-inducible *Pp*EC15 gene

Transgenic soybean plants expressing *Pp*EC15 were produced via Agrobacterium-mediated transformation using EHA105 cells, according to an established protocol (Luth et al., 2015). EHA105 cells containing pBAV154:*Pp*EC15:miniTurbo:V5, under a DEX-inducible promoter (Helm et al. 2019), were utilized for soybean transformation. 50 μM of DEX solution containing 0.02% Tween^®^20 was used to spray soybean leaves to induce cassette expression. All experiments were performed using the first or second fully expanded trifoliate 24 hours after induction.

### Protein extraction and immunoblot analysis

Total proteins were extracted as described previously (Padmanabhan et al., 2013) and separated on an SDS-PAGE gel, followed by immunoblotting using a horseradish peroxidase (HRP) system. Membranes were blocked with 5% non-fat milk in TBS-T (0.1% Tween-20 in Tris-buffered saline) before overnight incubation with indicated antibodies. After incubation with the appropriate HRP-conjugated secondary antibodies, bands were visualized by chemiluminescence (Pierce ECL). Primary anti-V5 (ChromoTek; Catalog number: v5ab) was used in a dilution 1:5000, and secondary HRP-conjugated Affinipure Goat Anti-Mouse was used in a dilution of 1:5000 with overnight incubation.

### Subcellular localization

The coding sequences of *GmPCRF* (Glyma.09G248300), *GmNAC83* (Glyma.18g301500), *GmDAHP* (Glyma.14g176600)*, Pp*EC15wt and mutant variants, without signal peptides and stop codons were cloned into pCR^TM^8/GW/TOPO^TM^ (Invitrogen) generating entry vector plasmids. Homemade vectors, 35S-ATG-GW-RFP, 35S-ATG-GW-mVenus or 35S-mVenus-GW, modified from pEarleyGate 100 (Earley et al., 2006), were used to generate *Pp*EC15-RFP, *Pp*EC15-mVenus or mVenus-*Pp*EC15 versions, respectively, using the Gateway cloning system (Invitrogen). *A. tumefaciens* GV3101 strains carrying the plasmid DNA were infiltrated into the leaves of *N. benthamiana* plants. Subcellular localization of the fusion proteins was detected using confocal microscopy (Zeiss Laser Scanning Microscopy 700) 2 days post-infiltration. Different fluorescent signals were excited with the following laser lines: mVenus (488 nm), 4′,6-diamidino-2-phenylindole (DAPI; 405 nm), and chlorophyll autofluorescence (555 nm). The emitted signals were detected using the following filters: mVenus (SP 555), DAPI (SP 460), and chlorophyll autofluorescence (LP 640).

### RNA isolation and reverse transcription real-time quantitative PCR

Approximately 100 mg of starting material was used for RNA isolation using the RNeasy Plant Mini Kit (Qiagen) for *N. benthamiana* or TRIzol reagent (Invitrogen) for soybean according to the manufacturer’s instructions. Two µg of RNA and SuperScript III First Strand kit (Invitrogen) for *N. benthamiana* or Maxima First Strand cDNA Synthesis kit with dsDNase (ThermoFisher) for soybean were used for cDNA synthesis. A quantitative real-time PCR assay was performed on a 7500 real-time PCR system (Applied Biosystems, CA, USA) using iTaq Universal SYBR Green Supermix (BioRad). *GmNREG3* (Glyma.08G211200) or *NbActin* (GenBank: AY594294.1) were chosen as internal controls to normalize the data. The transcription level was calculated using the 2^-ΔΔCT^ method (Livak and Schmittgen, 2001).

### Yeast constructs and two-hybrid screen

The Matchmaker LexA two-hybrid system (Clontech) was used for Y2H screening of an SBR-infected soybean cDNA library. *Pp*EC15, minus the signal peptide, was cloned into the pLexA vector to create a fusion with the DNA binding domain (BD). Approximately 2.6 x 10^6^ yeast transformants were screened on 2% SD/Gal/Raf/X-P-gal (-Ura/-His/-Trp/-Leu) following the manufacturer’s instructions (Clontech). Direct protein-protein interaction was confirmed by co-transformation of the respective plasmids into the yeast strain AH109 using the Matchmaker GAL4 Two-hybrid System (Clontech), followed by a selection of transformants on 2% SD (-Leu/-Trp) at 30 °C for 3 days and sub­sequent transfer to 2% SD/X-α-gal (-Leu/-Trp/-His) and 2% SD (-Leu/-Trp/-His/-Ade) to select for growth of interacting clones.

### Split-luciferase complementation

Split-luciferase assays were adapted from a previously published protocol (Chen et al. 2008). Briefly, leaf discs were collected from *N.benthamiana* 48 hours after Agrobacterium infiltration in 96-well plates containing 100 µL of water. Next, 75 µL of water was replaced with 25 µL of a 2 mM Luciferin (GoldBio; Catalog number: LUCNA-100) solution. The disks were incubated in the dark for 15 minutes and luminescence measurements were taken on a GloMax® Navigator Microplate Luminometer (Promega).

### Bimolecular fluorescence complementation

For BiFC constructs, *GmPCRF*, *GmDAHP*, *GmNAC83* and *Pp*EC15, minus the signal peptide, were amplified and cloned into the BiFC vectors pSPYNE-35S, pSPYCE-35S, phygll-SPYNE(R)l55 and pkanll-VYCE(R) (Walter et al., 2004; Waadt et al., 2008). Each protein was independently tagged with either cYFP or nYFP at the N- or C-terminus of the proteins of interest. *A. tumefaciens* GV3101 strains carrying the plasmids were infiltrated into *N. benthamiana* leaves. A BiFC signal was detected using a Zeiss LSM 700 confocal microscope 48 h after infiltration. YFP signal was excited at 488 nm and the emitted signal was collected using SP 555.

### *Pp*EC15 interactors cleavage assay

The digestion assay was performed with the purified native *Pp*EC15 proteins, wild type (wt) and double mutant (dm) using 2 µg of the interactor soybean proteins with 1 µg of *Pp*EC15 variants in PBS buffer (140 mM NaCl, 2.7 mM KCl, 10 mM Na_2_HPO_4_, 1.8 mM KH_2_PO_4_, pH 7.3) incubated at 37 °C for 16 hours. Proteins were visualized using Coomassie brilliant blue (CBB) staining.

### Virus-induced gene silencing

VIGS on soybean plants was performed according to a method previously described (Zhang et al., 2010; Whitham et al., 2016) with minor modifications. The coding sequences of *GmPCRF*, *GmNAC83*, and *GmDAHP* were used in the Sol Genomics Network VIGS design tool (https://vigs.solgenomics.net/) to select the VIGS fragments. Fragments amplified from soybean cDNA were cloned into RNA2 of the BPMV VIGS vector in the antisense orientation. BPMV::00 (EV) was used as a control. Inoculum was generated, stored at -20 °C, and used for the rub-inoculations as described previously (Zhang et al., 2010). The silencing efficiency was validated using RT-qPCR with primers targeting regions outside the insert region (Supplemental Table S3).

### Statistical analysis

One-way ANOVA and/or Student’s *t*-test were performed to calculate whether the differences between distributions of data were significant using PRISM v9.0 (GraphPad Software) and JMP PRO 16 software. A p-value of <0.05 was considered statistically significant.

## Acknowledgments

We acknowledge the W.M. Keck Metabolomics Research Laboratory (Office of Biotechnology, Iowa State University, Ames, IA) for providing analytical instrumentation, and we thank Ann M Perera, Lucas J Showman, and Matthew W Breitzman for their assistance and support. We thank Sarah Pottinger from the Innes laboratory for constructing the miniTurbo Gateway entry plasmid pBSDONR P4r-P2:miniTurbo. We also thank David Wright and Amber Testroet for their assistance with the transgenic soybean lines. Iowa State University is located on the ancestral territory of the Baxoje, or Ioway Nation; we are grateful to live, work, and play on these lands.

## Author Contributions

Designed the research: SAW, RWI, CSC, ASC, MQ, JWW, KA

Performed research: ASC, MQ, CM, HV, HD, AM, FC, MB, JM

Contributed reagents/materials/analysis tools: SAW, KFP, CSC, FC, KA, JWW

Analyzed the data: ASC, MQ, CM, HV, MB, JM, KA, FC

Wrote the paper: ASC, MB and SAW with contributions from all authors

## Commercial Endorsement Disclaimer

Mention of trade names or commercial products in this publication is solely for providing specific information and does not imply recommendation or endorsement by the U.S. Department of Agriculture (USDA).

## Equal Opportunity/Non-discrimination Statement

USDA is an equal opportunity provider and employer. The complete nondiscrimination policy can be found on the USDA website.

## Supplemental data

**Supplemental File 1. Supplemental Materials and Methods.**

**Supplemental Table S1. MiniTurbo-mediated proximity labeling.**

**Supplemental Table S2. Proximity labeling-identified *Nicotiana benthamiana Pp*EC15 predicted cleavage motifs.**

**Supplemental Table S3. Primers used in this study.**

**Supplemental Figure S1. *Pp*EC15 cleavage specificity.**

**Supplemental Figure S2. flg22-elicited apoplastic ROS production in *N. benthamiana* transiently expressing *Pp*EC15wt.**

**Supplemental Figure S3. *Pp*EC15 delivered by *P. fluorescens* expressing a *P. syringae* type III secretion system (TTSS) is able to suppress basal resistance in *N. benthamiana*, as indicated by the hypersensitive response (HR) challenge assay.**

**Supplemental Figure S4. Positive and negative controls for bimolecular fluorescence complementation (BiFC) experiment.**

**Supplemental Figure S5. *Pp*EC15-interacting protein enrichment by miniTurbo-based biotin labeling.**

**Supplemental Figure S6. *Pp*EC15-DAHP subcellular colocalization.**

**Supplemental Figure S7. *Pp*EC15-soybean interactors cleavage assay.**

