## Supplemental Figures for "A soybean rust effector protease suppresses host immunity and cleaves a 3-deoxy-7-phosphoheptulonate synthase"

**A**

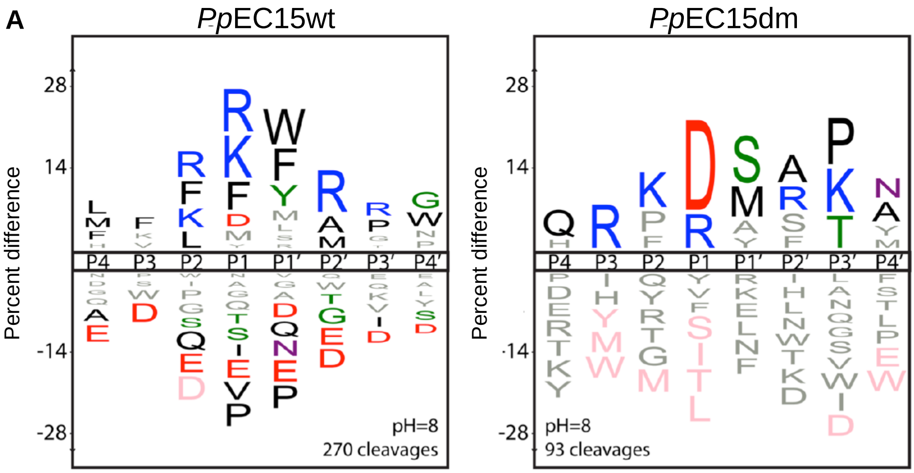


**B**

| Sequence | Ion Precursor Intensities | | | | | | Additional cleavages |
|---|---|---|---|---|---|---|---|
| | wt | | | dm | | | |
| | 60' | 240' | 1200' | 60' | 240' | 1200' | |
| YKRF/MAHW | 53668 | 22092 | 96043 | 0 | 0 | 0 | P1', P3 |
| MFRK/YPIM | 12501 | 38450 | 928860 | 0 | 0 | 0 | |
| SLYR/MIRQ | 0 | 15733 | 50267 | 0 | 0 | 13147 | |
| MHRN/WRAQ | 10450 | 29486 | 169759 | 0 | 0 | 0 | |
| HVKL/FRFN | 9317 | 14836 | 195553 | 0 | 0 | 0 | P1', P2' |
| PTVN/KQLR | 0 | 6476 | 20154 | 0 | 0 | 0 | P1', P2' |

**Supplemental Figure S1. *Pp*EC15 cleavage specificity. A.** Motif analysis of time-dependent substrate specificity profiles for *Pp*EC15wt and *Pp*EC15dm is plotted as fold enrichment at each position P4 to P4'. Residues above the line are favored; residues below the line are disfavored. Statistically significant residues (p <0.05) are colored by physicochemical properties (black: hydrophobic; blue: basic; red: acidic). Residues in gray were not statistically significant, but they were observed, and residues in pink are absolutely disfavored (never observed in a cleavage). Letter height corresponds to the fold enrichment value**. B.** Curated list of selective cleavages from the full MSP-MS library (228 14-mer proprietary peptide substrates). Each cleavage is represented as an 8-mer peptide from P4 to P4', where the P1-P1' position represents the scissile bond. Label-free quantitation of precursor ion abundance from the MS/MS spectra is reported for each cleavage at the P1-P1' position; the last column reports additional cleavage events at other positions when detected.

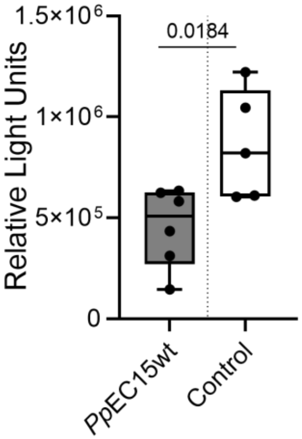


**Supplemental Figure S2. flg22-elicited apoplastic ROS production in *N. benthamiana* transiently expressing *Pp*EC15wt.** Data are represented as box plots indicating the 25%–75% interquartile range, split by a median line. Whiskers represent maximum and minimum values. Statistically significant values ($p < 0.05$) are determined by a *t*-test using GraphPad Prism 9.0. The experiment was conducted at least three times with similar results.

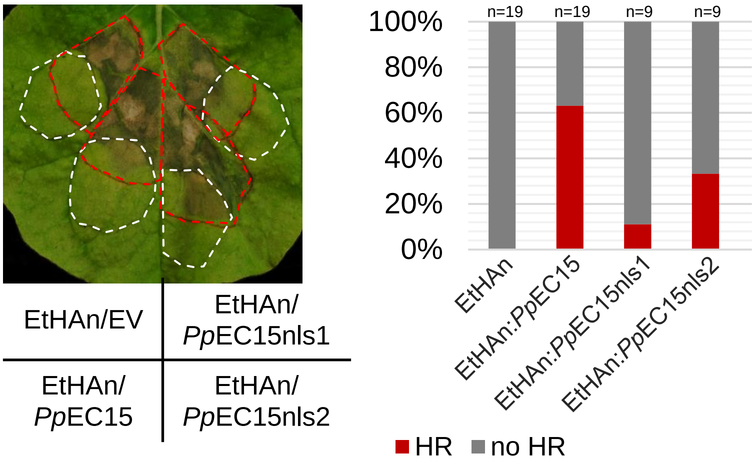


**Supplemental Figure S3. *Pp*EC15 delivered by *P. fluorescens* expressing a *P. syringae* type III secretion system (TTSS) is able to suppress basal resistance in *N. benthamiana*, as indicated by the hypersensitive response (HR) challenge assay.** *N. benthamiana* leaves were infiltrated with *P. fluorescens* (EtHAn) carrying an empty vector or expressing *Pp*EC15wt or mutant versions, and then challenged 6 h later by overlapping inoculation with *P. syringae* pv. tomato DC3000 at $2 \times 10^7$ CFU $ml^{-1}$. The white and red dotted lines indicate the area infiltrated with *P. fluorescens* (EtHan) and *Pto*DC3000, respectively. Leaves were photographed 24 h after inoculation with *Pto*DC3000. Percentage of leaves in which an HR was observed in the overlapping area are shown in the graph.

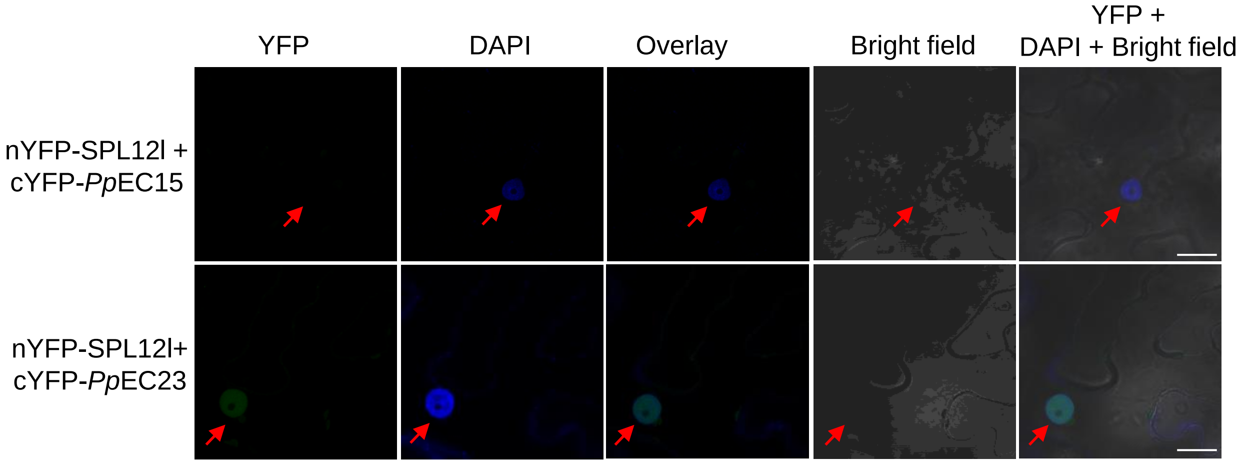


**Supplemental Figure S4. Positive and negative controls for bimolecular fluorescence complementation (BiFC) experiment.** BiFC assay showing the negative (SPL12+*Pp*EC15) and positive (SPL12l+*Pp*EC23) controls *in planta*. DAPI signal was used as a nuclear marker. Arrows indicate nuclei. Representative images are shown (n=20). Scale bars = 20 µm.

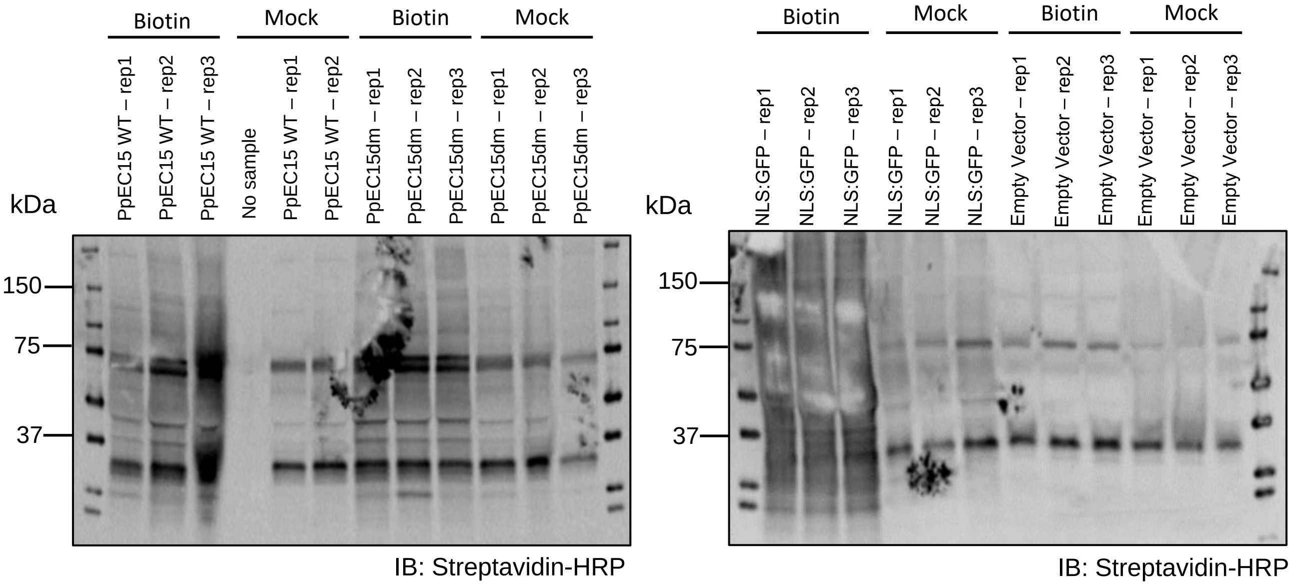


**Supplemental Figure S5. *Pp*EC15-interacting protein enrichment by miniTurbo-based biotin labeling.** Analysis of biotinylated proteins from *N. benthamiana* leaves 2 hours after infiltration with 200 µM biotin solution or mock solution. Biotinylated proteins of three replicates (Rep 1-3) were detected using Streptavidin-HRP conjugated antibody 1:5000 dilution.

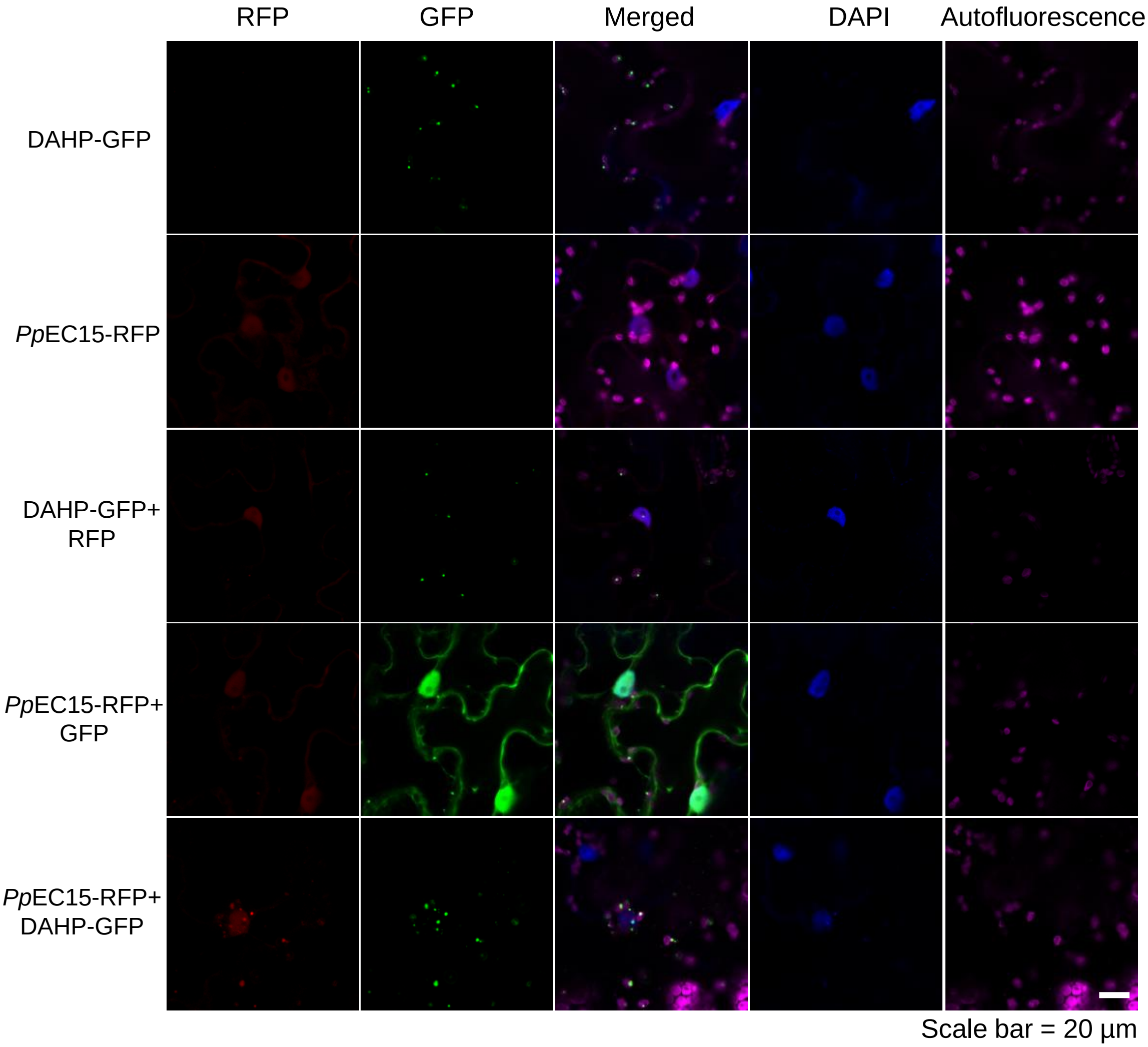


**Supplemental Figure S6. *Pp*EC15-DAHP subcellular co-localization.** Subcellular localization of RFP-tagged *Pp*EC15 and GFP-tagged DAHP proteins. Red fluorescence indicates the localization of *Pp*EC15, and green fluorescence indicates the localization of DAHP in *N. benthamiana* epidermal cells. DAPI signal was used as a nuclear marker. Representative images are shown (n=20). Scale bars = 20 µm. Photographs were taken at 48 hpi using confocal microscopy.

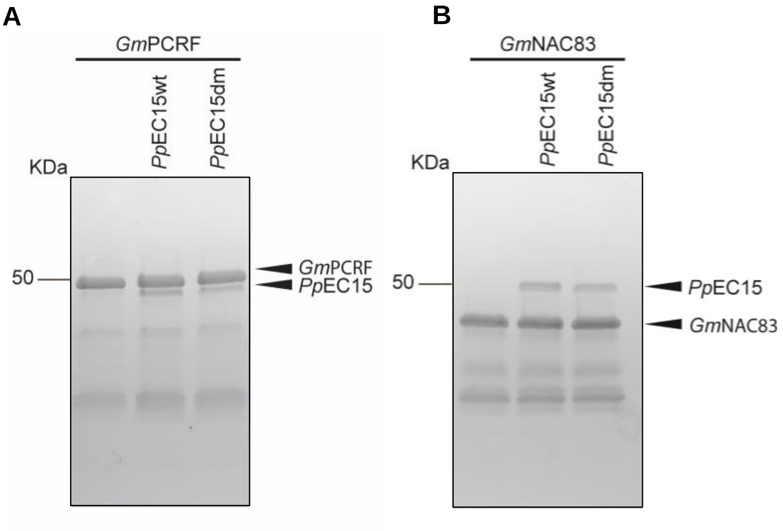


**Supplemental Figure S7. *Pp*EC15 interactors cleavage assay.** SDS-PAGE demonstrating the cleavage assay performed with *Pp*EC15wt and *Pp*EC15dm with *Gm*PCRF **(A)** and *Gm*NAC83 **(B)**. 1 µg of recombinant *Pp*EC15 protein and 2 µg of the recombinant soybean proteins were incubated at 37 °C for 16 hours. Proteins were visualized using Coomassie brilliant blue (CBB) staining.
