## Supplemental File 1 for "A soybean rust effector protease suppresses host immunity and cleaves a 3-deoxy-7-phosphoheptulonate synthase"

### Supplemental Materials and Methods

#### *Pp*EC15 peptide cleavage specificity assay

To determine the cleavage specificity of *Pp*EC15wt and *Pp*EC15dm, samples were used at a final concentration of 500 nM in a buffer containing 50 mM Tris (pH 8.0) and 100 mM NaCl. Samples and matched no-enzyme controls were incubated with a library of 228 14-mer synthetic peptide substrates (O'Donoghue et al. 2015). Aliquots were collected and acid quenched at 60, 240, and 1200 min time points. All samples were desalted with C18 desalting tips, lyophilized, and rehydrated in 0.1% formic acid. Peptide sequencing by LC-MS/MS was performed on an LTQ- Orbitrap XL mass spectrometer (Thermo) equipped with a nanoACQUITY (Waters) ultra-performance LC (UPLC) system and an EASYSpray ion source (Thermo). Reversed-phase chromatography was carried out with an EASYSpray PepMap C18 column (Thermo, ES800; 3-m bead size, 75 m by 150 mm). LC was performed at a 600-nl/min flow rate during sample loading for 14 min and then at a 300-nl/min flow rate for peptide separation over 65 min with a linear gradient of 2 to 50% (vol/vol) acetonitrile in 0.1% formic acid. Peptide fragmentation was performed by collision-induced dissociation (CID) on the six most intense precursor ions, with a minimum of 1,000 counts, with an isolation width of 2.0 m/z, and a minimum normalized collision energy of 25. MS peak lists were generated with MSConvert, and data were searched against the 228-member peptide library with Protein Prospector software, v.5.16.1 (http://prospector.ucsf.edu/prospector/mshome.htm, UCSF), and tolerances of 20 ppm for parent ions and 0.8 Da for fragment ions. All cleavages were allowed in the search by designating no enzyme specificity. The following variable modifications were used: amino acid (proline, tryptophan, and tyrosine) oxidation and N-terminal pyroglutamate conversion from glutamine. Protein Prospector score thresholds were selected with a minimum protein score of 15 and a minimum peptide score of 15. Maximum expectation values of 0.01 and 0.05 were selected for protein and peptide matches, respectively. Peptides corresponding to cleavage products in the 228-member library were imported into iceLogo software v.1.2 to generate substrate specificity profiles. Octapeptides corresponding to P4-P4' were used as the positive data set, and octapeptides corresponding to all possible cleavages in the library (n=2,964) were used as the negative data set. Heatmaps were created in the R environment using the gplot and RColorBrewer packages.

#### Proximity labeling constructs

*Pp*EC15wt and *Pp*EC15dm, without the signal peptides, and NLS-GFP coding sequence were fused to the N-terminus of miniTurbo by MultiSite gateway cloning. Briefly, *Pp*EC15wt, *Pp*EC15dm, and NLS:GFP were cloned into pBSDONR P1-P4 gateway-compatible entry vector by BP clonase reaction (Invitrogen), and the clones were sequence-verified. pBSDONR P1-P4:*Pp*EC15, pBSDONR P1-P4:*Pp*EC15dm, pBSDONR P1-P4:NLS:GFP were then combined with pBSDONR P4r-P2:miniTurbo and pEG100 (Earley et al., 2006) or pBAV154 (Helm et al. 2019) destination vectors in an LR clonase reaction (Invitrogen).

#### Preparation of biotin proximity labeled protein samples

Fully expanded 4-5 weeks-old *N. benthamiana* leaves were infiltrated with *A. tumefaciens* GV3101 containing either pEG100::*Pp*EC15wt:miniTurbo, pEG100::*Pp*EC15dm:miniTurbo or pEG100::NLS:GFP:miniTurbo constructs at $OD_{600}$=1.0. Thirty-six hours post-infiltration, a 200 µM biotin solution was infiltrated in the same leaf areas and 2 h later, samples were collected and stored at -80 °C until use. Protein extraction and preparations were performed according to a recently published proximity labeling protocol optimized for plants (Zhang et al. 2020; Zhang et al. 2019). In brief, approximately 35 mg of plant tissue was ground in RIPA lysis buffer (50 mM Tris-HCl [pH 7.5], 500 mM NaCl, 1 mM EDTA, 1% NP40 [v/v], 0.1% SDS [w/v], 0.5% sodium deoxycholate [w/v], 1 mM DTT, 1 tablet of protease inhibitor cocktail). Total protein was collected by centrifugation, and removal of free biotin was performed by using Zeba™ Spin Desalting Columns, 7K MWCO, 10 mL (Catalog number: 89893) according to the manufacturer's instructions. Quantification of the desalted protein extracts was performed using Bradford assay, and 1 mg of protein was used for the enrichment of the biotinylated proteins by utilizing Dynabeads™ MyOne™ Streptavidin C1 (Catalog number: 65001) according to the manufacturer's instructions.

**Identification of biotinylated peptides by mass-spectrometry**

Biotinylated protein samples from *N. benthamiana* were reduced in 2 mM TCEP (tris(2-carboxyethyl)phosphine) and digested into peptides at 37 °C overnight in 0.2 µg trypsin plus 0.1 µg Lys-C. Digested peptides were alkylated in 50 mM iodoacetamide (IAM). Peptides were further desalted using SepPack C18 columns (Waters). Tandem Mass Tag (TMT, Thermo Scientific) labeling was performed on purified peptides from each sample, as previously reported (Song et al., 2020). TMT labeling reaction was stopped using 5% hydroxylamine, and the quenched samples were then pooled. Chromatography was performed on a Thermo UltiMate 3000 UHPLC RSLCnano. Peptides were desalted and concentrated on a PepMap100 trap column (300 µM i.d. x 5 mm, 5 µm C18, 100 Å µ-Precolumn, Thermo Scientific) at a flow rate of 10 µL $min^{-1}$. Sample separation was performed on a 200 cm Micro-Pillar Array Column (µ-PAC, Pharmafluidics) with a flow rate of ~300 nL $min^{-1}$ over a 150 min reverse phase gradient (80% ACN in 0.1% FA from 1% to 15% over 5 min, 15% to 20.8% over 20 min, from 20.8% to 43.8% over 80 min, from 43.8% to 99.0% in 11 min and kept at 99.0% for 5 min). Eluted peptides were analyzed using a Thermo Scientific Q-Exactive Plus high-resolution quadrupole Orbitrap mass spectrometer, which was directly coupled to the UHPLC. Data-dependent acquisition was obtained using Xcalibur 4.0 software in positive ion mode with a spray voltage of 2.3 kV and a capillary temperature of 275 °C and an RF of 60. MS1 spectra were measured at a resolution of 70,000, an automatic gain control (AGC) of 3e6 with a maximum ion time of 100 ms and a mass range of 400-2000 m/z. Up to 15 MS2 were triggered at a resolution of 35,000. A fixed first mass of 115 m/z. An AGC of 1e5 with a maximum ion time of 50 ms, a normalized collision energy of 33, and an isolation window of 1.3 m/z were used. Charge exclusion was set to unassigned, 1, 5–8, and >8. MS1 that triggered MS2 scans were dynamically excluded for 25 s. Raw data were analyzed using MaxQuant version 1.6.14 (Cox and Mann, 2008). Spectra were searched using the Andromeda search engine (Cox et al., 2011) against *N. benthamiana* annotation v1.0.1 (Bombarely et al., 2012). The proteome files were complemented with reverse decoy sequences and common contaminants by MaxQuant. Carbamidomethyl cysteine was set as a fixed modification, while methionine oxidation and protein N-terminal acetylation were set as variable modifications. The sample type was set to “Reporter Ion MS2" with “TMT11plex” selected for both lysine and N-termini. Digestion parameters were set to “specific” and “Trypsin/P;LysC”. Up to two missed cleavages were allowed. A false discovery rate, calculated in MaxQuant using a target-decoy strategy (Elias and Gygi, 2007), less than 0.01 at both the peptide spectral match and protein identification level was required. The match between runs feature of MaxQuant was not utilized. Differential protein enrichment was assessed using the PoissonSeq R package (Li et al., 2012), implemented in our TMT-NEAT proteomics analysis pipeline (Clark et al., 2021). A q-value < 0.1 was used as the cutoff to define *Pp*EC15-enriched proximal proteins.

**Gas chromatography-mass spectrometry non-targeted metabolomic analysis in VIGS soybean**

Sample preparation was conducted using a modified version of the methanolic extraction and sample preparation methods established by (A et al., 2005). Approximately 50 mg of each sample was spiked with internal standard 10 µg ribitol (1 mg/mL in water) (Sigma-Aldrich CO., St. Louis, MO). 0.9 mL of 80% LC-MS grade methanol with 20% LC-MS grade water (Fisher Scientific, Waltham, MA) was added to the samples. Samples were incubated for 10 min at room temperature, vortexed for 30 sec, and placed into an ice-cold sonication water bath for 10 min. Samples were vortexed for 30 sec and centrifuged for 10 min at 16,000 x g. Supernatants were recovered, and the remaining insoluble pellets were re-extracted with an additional volume of 0.9 mL of 80% methanol, and the supernatant extracts were pooled. Six hundred µL of the combined extracts were dried using a speed-vac concentrator for 10 h prior to derivation (Koek et al., 2006). Samples were derivatized with 50 µL of methoxyamine hydrochloride (20 mg/mL in pyridine) initially added to the dried extracts, followed by a 1.5 h incubation at 30 °C. Subsequently, timethylsilylation (TMS) was performed by the addition of 70 µL of bis-trimethylsilyl trifluoroacetamide with 1% Trimethylchlorosilane (BSTFA + 1% TMCS) for 30 min at 60 °C. Derivatized samples were analyzed by GC-MS with an Agilent 6890 gas chromatograph coupled to a model 5973 Mass Selective Detector (Agilent Technologies, Santa Clara, CA). The column used was HP-5MSI 5% Phenyl Methyl Silox with 30 m × 250µM × 0.25 µm film thickness (Agilent Technologies). One microliter of sample was injected with the inlet operating in splitless mode and held at a constant temperature of 280 °C. The oven temperature was programmed as follows: an initial temperature of 70 °C was increased to 320 °C. The MS transfer line was held at 280 °C. MS detection was performed using electron ionization at 70 eV and source temperature and quadrupole temperature were set at 230 °C and 150 °C, respectively. The mass data were collected in the range from m/z 40 to m/z 800. Identification and quantification were conducted using AMDIS (Automated Mass spectral Deconvolution and Identification System, National Institute of Standards and Technology (Gaithersburg, MD)) with a manually curated retention indexed GC-MS library with additional identification performed using the NIST17[b2] and Wiley 11 GC-MS spectral library (Agilent Technologies, Santa Clara, CA). Final quantification was calculated by integrating the corresponding peak areas relative to the area of the internal standards. Raw data were normalized to the amount of tissue used. Statistical evaluation of the non-targeted GC-MS data was conducted with the R-based statistical package MetaboAnalyst (Pang et al., 2021).

**DAHP homotetramer protein modeling**

The 3D structures of DAHP were modeled through Phyre and visualized using the PyMOL Molecular Graphics System. The homotetramer structure of *Gm*DAHP was predicted through Alphafold multimer (ColabFold: https://colab.research.google.com/github/sokrypton/ColabFold/blob/main/AlphaFold2.ipynb) using 6 recycles per model. The best model among five was selected (pTM = 0.833) and aligned with the homotetramer of DAHP from *Corynebacterium glutamicum* in complex with chorismate mutase (PDB: 5HUD). The structural alignment was visualized using PyMOL.
